# Requirements for mammalian promoters to decode transcription factor dynamics

**DOI:** 10.1101/2021.10.12.464037

**Authors:** Enoch B. Antwi, Yassine Marrakchi, Özgün Çiçek, Thomas Brox, Barbara Di Ventura

**Affiliations:** Heidelberg Biosciences International Graduate School (HBIGS), University of Heidelberg, Heidelberg, Germany; Centers for Biological Signalling Studies BIOSS and CIBSS, Albert-Ludwigs-University, Freiburg, Germany; Faculty of Biology, Institute of Biology II, Albert-Ludwigs-University, Freiburg, Germany; Department of Computer Science, Albert-Ludwigs-University, Freiburg, Germany; BrainLinks-BrainTools, Albert-Ludwigs-University, Freiburg, Germany

**Keywords:** Transcription factor, dynamics, gene expression, optogenetics, mathematical modeling

## Abstract

In response to different stimuli many transcription factors (TFs) display different activation dynamics that trigger the expression of specific sets of target genes, suggesting that promoters have a way to decode them. Combining optogenetics, deep learning-based image analysis and mathematical modeling, we find that decoding of TF dynamics occurs only when the coupling between TF binding and transcription pre-initiation complex formation is inefficient and that the ability of a promoter to decode TF dynamics gets amplified by inefficient translation initiation. Furthermore, we propose a theoretical mechanism based on phase separation that would allow a promoter to be activated better by pulsatile than sustained TF signals. These results provide an understanding on how TF dynamics are decoded in mammalian cells, which is important to develop optimal strategies to counteract disease conditions, and suggest ways to achieve multiplexing in synthetic pathways.

## Introduction

Gene expression is a tightly regulated, complex biological process that turns a specific DNA sequence, the gene, into RNA or protein. It comprises several steps, among which are transcription, mRNA translation, and protein folding and degradation (Buccitelli and Selbach, 2020). In eukaryotes, transcription of protein-coding genes is carried out by RNA polymerase II (Pol II) and typically initiates at the transcription start site (TSS) found at the 5’ end of a gene within the core promoter. This is the location at which the general transcription machinery ––constituted by Pol II and its associated general transcription factors (GTFs) (Hampsey, 1998)–– assemble, forming the pre-initiation complex (PIC) (Haberle and Stark, 2018). The core promoter has specific sequence motifs, which are known to recruit GTFs to mediate PIC assembly. The two Pol II core promoter motifs capable of nucleating the PIC are the TATA box and the Initiator element (Inr) (Roy and Singer, 2015). Core promoters usually have low basal activity and are regulated by distal DNA elements –– the enhancers–– as well as chromatin state (Kornberg and Lorch, 2020; Zabidi and Stark, 2016). Enhancers are the genomic regions at which transcription factors (TFs) and co-factors are recruited (Kadonaga, 2004; Lambert et al., 2018). TFs bind specific DNA sequences called response elements (REs) (Zhu and Huq, 2011). Individual TFs often control a multitude of genes (Purvis and Lahav, 2013), and they act either as repressors or activators. DNA loop formation brings the TF-bound RE(s) and the core promoter into close proximity, allowing the TF to recruit the GTFs (Bartman et al., 2016; Petrascheck et al., 2005).

Specialization of TFs for certain targets is one of the mechanisms used by cells to start different gene expression programs under specific conditions. The human genome codes for approximately 1,600 TFs (Lambert et al., 2018). Likely the necessity to maintain a manageable genome size led to the evolution of other strategies consenting cells to re-use the same TF in multiple ways. One strategy consists in post-translationally modifying TFs to modulate their stability, localization as well as affinity for the DNA, co-activators and/or the GTFs (Moore et al., 2011; Tootle and Rebay, 2005). In the past decade, another cellular strategy emerged as an effective way of achieving multiplexing: controlling TF dynamics, that is, the time-resolved activity of the TF (Behar and Hoffmann, 2010; Behar et al., 2013; Purvis and Lahav, 2013). In particular, several studies have shown that TFs accumulate in the nucleus –where they are active– either in a single, sustained pulse, or in repeated pulses of distinct frequencies and amplitudes depending on the stimulus sensed by the cells (Muta et al., 2019; Purvis and Lahav, 2013). p53, for instance, in response to UV irradiation, was shown to accumulate in the nucleus in a single pulse, with amplitude proportional to the UV dose, and in pulses of fixed amplitude in response to *γ*-radiation (Batchelor et al., 2011). Artificially turning the natural p53 pulsatile dynamics in response to *γ*-radiation into a single sustained pulse increased the frequency of cells going into senescence instead of recovering from the DNA damage, suggesting that TF dynamics directly influence cell fate decisions (Purvis et al., 2012).

If TF dynamics dictate which genes are activated, then promoters have a way to decode them. TF dynamics decoding at the level of the promoter has been so far studied in the yeast *Saccharomyces cerevisiae* (Hansen and O’Shea, 2013) and the filamentous fungus *Neurospora crassa* (Li et al., 2018). A systematic analysis of the regulatory elements necessary to render a mammalian promoter sensitive to TF dynamics is missing. Moreover, the role played by translation initiation efficiency in the decoding process remains unclear.

In this paper, we apply optogenetic perturbations to control the nuclear localization of a synthetic TF, and generate defined TF dynamics. We then study the ability of these TF dynamics to activate a library of synthetic promoters built of well-defined and characterized parts. Furthermore, we investigate the effect of translation initiation rates in transmitting TF dynamics to downstream gene expression programs. Combining experiments and mathematical modelling, we show that different TF dynamics are distinguished by promoters characterized by inefficient coupling between TF binding and PIC formation and stabilization, and that inefficient translation of the mRNA amplifies the decoding process achieved at the promoter level. Using the mathematical model, we explore a mechanism that would allow a promoter to filter out sustained signal while responding to TF pulses.

## Results

### Different TF dynamics can be imposed using an engineered light-responsive synthetic TF

We constructed a synthetic TF (synTF) fusing to well-characterized *E. coli* repressor protein LexA (Howard-Flanders et al., 1966) three copies of residues 436-447 of the herpes simplex virus type 1 transcription factor VP16 (VP48) (Triezenberg et al., 1988), the fluorescent protein mCherry and LINuS, an optogenetic tool consisting of a light-inducible nuclear localization signal (NLS) that allows accumulating synTF in the nucleus or the cytosol upon blue light illumination and incubation in the dark, respectively (Niopek et al., 2014) (Figures 1A and S1A). By selecting the appropriate light regime, we can generate different synTF dynamics; here we opted for a single sustained pulse (we refer to this as sustained activation), pulses with ~30-min period (designated 15-15 pulses) and pulses with ~45-min period (designated 15-30 pulses) (Figure S1B). synTF transcriptional activity is quantified at the reporter nascent RNA (Figure S1C) and protein (Figure S1D) levels. To capture differences due solely to TF dynamics, and not cumulative TF levels, the experiments are assigned different durations, but they all comprise the same five-hour waiting time at the end of the last activation phase to allow for the maturation of the iRFP670 reporter protein (Shcherbakova and Verkhusha, 2013) (Figure 1B). The workflow starts with a transient transfection step, explicitly chosen to obtain information about the sensitivity of the promoters towards synTF amplitude, followed by the application of the dynamics, and the automated quantification of the time-lapse microscopy images (Figure 1C). The image processing pipeline is based on a neuronal network developed by us for the specific task of segmentation in the absence of dedicated nuclear and plasma membrane fluorescent markers (Çiçek et al., 2020). The data are finally clustered into bins based on nuclear synTF level (mCherry signal) at t=0 for fair comparison of different dynamics and promoters.

**Figure 1.**
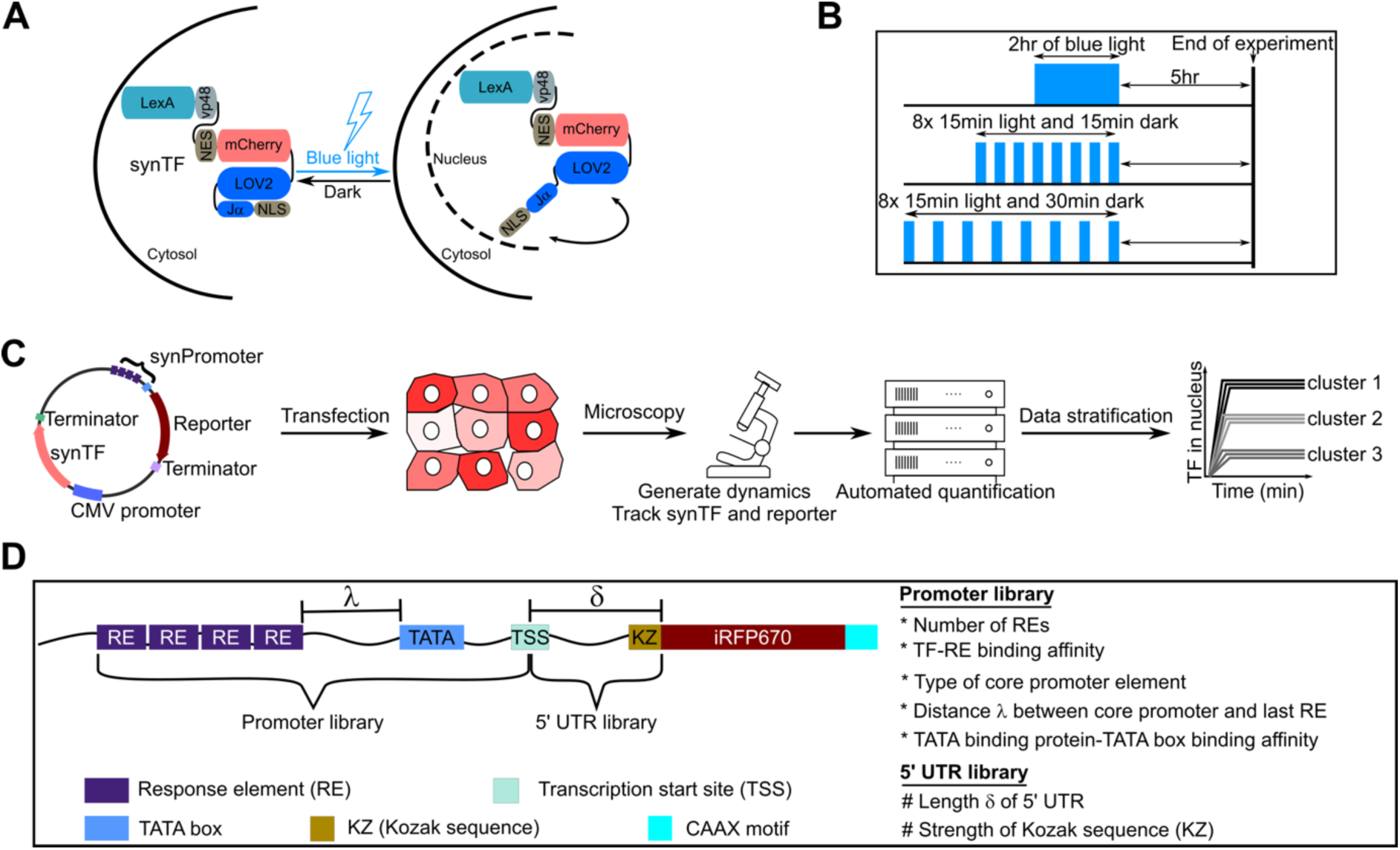
Design of molecular components and experimental setup used to study how transcription factor dynamics are decoded by mammalian gene regulatory elements. (A) Schematic representation of the expected behavior of the synthetic transcription factor (synTF) in darkness (left) and during blue light illumination (right). Blue light leads to the exposure of the NLS, which, in turn, leads to the nuclear import of synTF. LexA, full-length bacterial repressor used as DNA binding domain (DBD) within synTF. VP48, 3x residues 436-47 of the VP16 transactivation domain (TAD). NES, nuclear export signal. (B) Schematic representation of the illumination schemes used in this study to achieve similar cumulative TF levels (area under the curve) for the different TF dynamics. (C) Schematic overview of the experimental setup. (D) Schematic representation of the reporter libraries used in this study.

To understand the contribution of specific regulatory elements to the TF dynamics decoding process, we built a library of reporter constructs made of well-characterized DNA elements. As the gene regulatory elements, we focused on TF binding site (RE), core promoter, and 5’UTR. We created two types of libraries: one in which the promoter varies, and one in which the 5’UTR does (Figure 1D and Table S1). The TF and the reporter gene are encoded on the same plasmid to ensure that lack of iRFP670 signal in individual cells be not due to the absence of the reporter gene (Figure 1C).

### Mammalian promoters can decode TF dynamics

We started analyzing the first promoter in the library, promoter p1, which is characterized by four repeats of a strong RE and a strong TATA box (Table S1). We imposed to the cells the three different light regimes for sustained and pulsatile dynamics, and measured synTF nuclear concentration (Figure 2A), mean reporter nascent RNA (Figure 2B) and protein (Figure 2C) levels over time. Nascent RNA visualization was performed only during the blue light illumination phases not to activate the LOV domain within LINuS, which would result in synTF nuclear localization and promoter binding. The RNA transcription status during the dark phases can be deduced from the first image acquired in each activation phase. All three dynamics elicited a robust activation of promoter p1. The RNA data showed that there is no transcriptional shutdown during the dark phases for the pulses (Figure 2B, middle and right panels). When comparing the mean reporter nascent RNA levels per cell per minute, we did not find any difference between conditions (Figure 2D), corroborating the conclusion that promoter p1 does not distinguish dynamics. Interestingly, at the protein level, both pulsatile dynamics triggered a higher response than sustained activation (Figure 2E). Promoter p1 does not respond significantly to synTF amplitude (Figure 2F). When we tested a version of promoter p1 with the Inr in place of the TATA box as core promoter motif, we found only background levels of the reporter protein for sustained synTF dynamics. Hence all promoters we discuss have the TATA box as core promoter motif.

**Figure 2.**
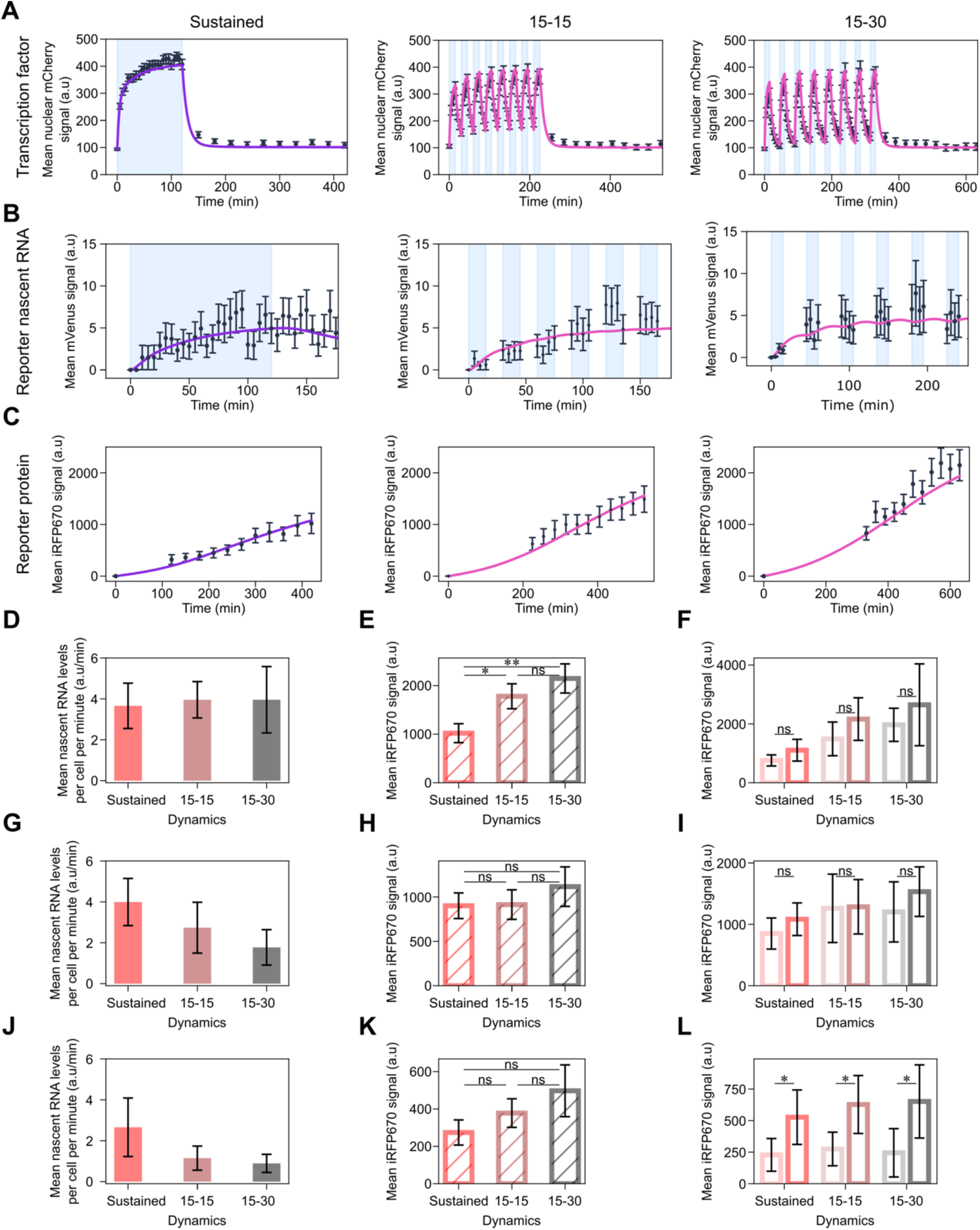
A promoter with strong RE and TATA box does not distinguish TF dynamics. (A-C) Quantification of synTF nuclear translocation (A), mean reporter nascent RNA (B) and protein (C) levels over time for the indicated TF dynamics in combination with promoter p1. Light blue shadowing, blue light illumination phase. Together with the experimental data (black dots), fitted (violet line; for sustained dynamics) and simulated (pink line; for both pulsatile dynamics) values are shown. The mathematical model used is shown in Figure 4 and the equations are described in the Supplememtary Text. (D,G,J) Quantification of mean nascent RNA per cell per minute for the indicated synTF dynamics in combination with promoters p1 (D), p2 (G) and p3 (J). (E,H,K) Quantification of reporter protein expression levels at the end of the experiment for the indicated synTF dynamics in combination with promoters p1 (E), p2 (H) and p3 (K). (F,I,L) Quantification of reporter protein expression levels at the end of the experiment for high (dark colors) or low (light colors) synTF amplitudes for the indicated synTF dynamics in combination with promoters p1 (F), p2 (I) and p3 (L). (E,F,H,I,K,L) P-values were calculated with the Welch’s t-test. ns, non signficant (P > 0.05). (E) *, P-value = 0.0244; **, P-value = 0.003. (L) *, P-value (from left to right) = 0.02552, 0.01285, 0.03517. Data represent mean ± s.e.m. of at least n=20 individual cells, imaged on at least n=3 biologically independent experiments.

Next, we analyzed promoters p2 and p3 (Table S1), which are characterized by either the same strong RE as promoter p1, but a weak TATA box (p2) or the same strong TATA box, but a weak RE (p3). We again imposed the three light regimes and measured synTF nuclear concentration (Figure S2A), mean reporter nascent RNA (Figures S2B and S2D) and protein (Figures S2C and S2E) levels over time. Interestingly, we observed a refractory response for promoter p2 under sustained activation, reflected in the decrease of the mean nascent RNA levels during the activation phase (Figure S2B, left panel). Differently than for promoter p1, we found lower mean reporter nascent RNA levels per cell per minute for the pulsatile than for the sustained dynamics for both promoters (Figures 2G and 2J). Thus, these promoters are able to distinguish TF dynamics. At the protein level, however, we found no significant difference at the end of the experiment (Figures 2H and 2K). While promoter p2 is insensitive to synTF amplitude (Figure 2I), promoter p3 is (Figure 2L). This is in line with the fact that promoter p3 features a weak RE, which requires a higher synTF amplitude to achieve full promoter activation.

Finally, we analyzed promoter p4, which combines a weak RE (four repeats thereof) with a weak TATA box (Table S1, and Figures 3A-3C). For promoter p4, refractoriness was more prominent than for promoter p2 (compare Figures S2B and 3B, left panels). Despite being overall a weaker promoter than p1-p3, p4 is the best at distinguishing synTF dynamics. Indeed, the two pulsatile dynamics elicited only a mild activation of promoter p4, and, in this case, this is seen both, at the reporter RNA (Figure 3D) and protein (Figure 3E) levels. Interestingly, the difference between the responses of promoter p4 to two different synTF amplitudes appears not statistically significant for sustained and 15-30 pulses (Figure 3F). Nonetheless, since promoter p4 is characterized by a weak RE, we consider this promoter sensitive to amplitude. Lack of statistical significance is justified by the noisy nature of this promoter (Figure S3). The more stochastic nature of promoter p4 compared to promoters p1-p3 is justified by its strong refractory behavior, because exiting the refractory state is a random process (Harper et al., 2011).

**Figure 3.**
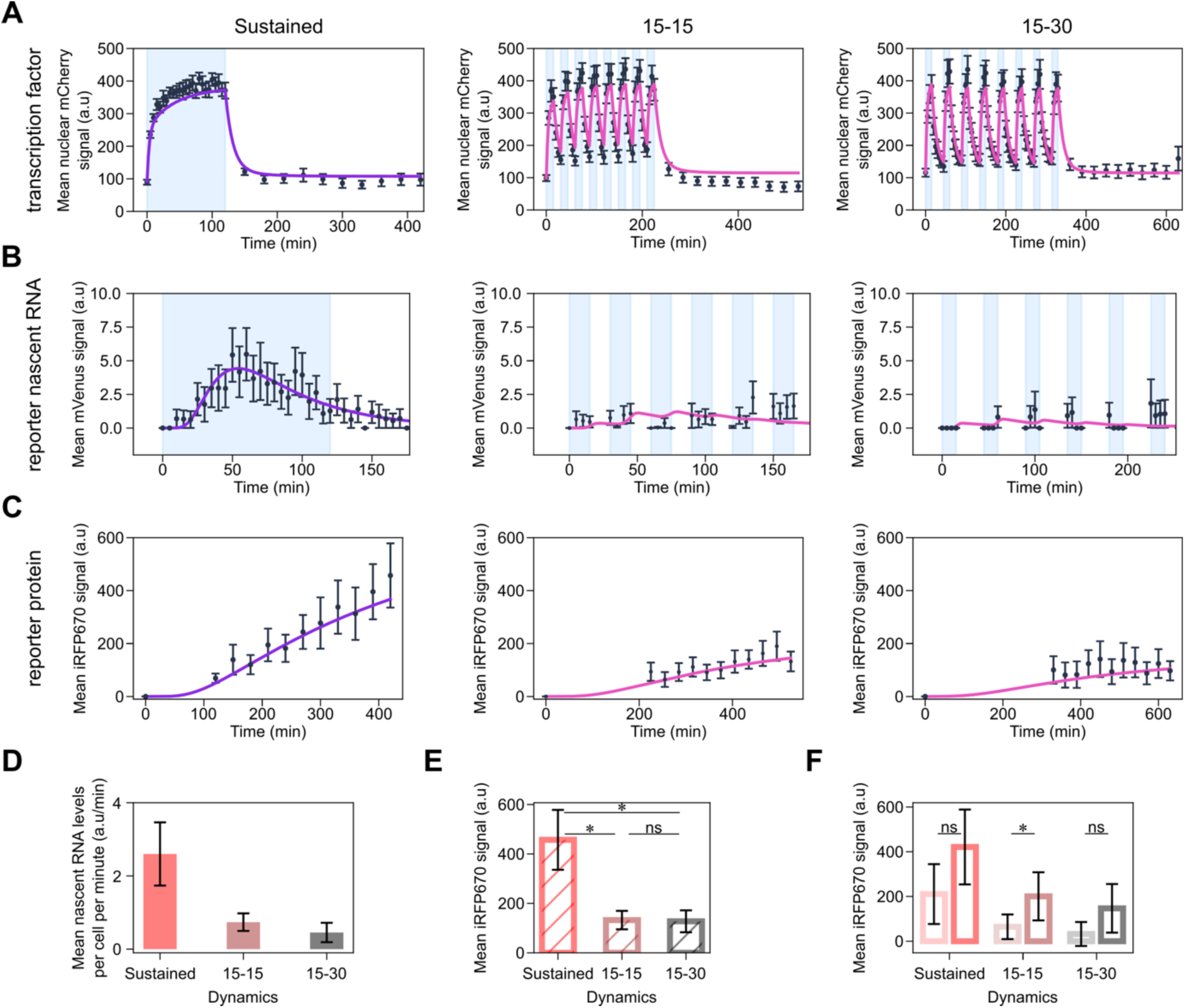
A promoter with weak RE and TATA box distinguishes TF dynamics. (A-C) Quantification of synTF nuclear translocation (A), mean reporter nascent RNA (B) and protein (C) levels over time for the indicated TF dynamics. Light blue shadowing, blue light illumination phase. Together with the experimental data (black dots), fitted (violet line; for sustained dynamics) and simulated (pink line; for both pulsatile dynamics) values are shown. The mathematical model used is shown in Figure 4 and the equations are described in the Supplememtary Text. (D) Quantification of mean nascent RNA per cell per minute for the indicated synTF dynamics. (E) Quantification of mean reporter protein expression levels at the end of the experiment for the indicated synTF dynamics. (F) Quantification of mean reporter protein expression levels at the end of the experiment for high (dark colors) or low (light colors) synTF amplitudes for the indicated synTF dynamics. (E,F) P-values were calculated with the Welch’s t-test. ns, non signficant (P > 0.05). (E) *, P-value (from left to right) = 0.0217; 0.0212. (F) *, P-value = 0.03292. (A-F) Data represent mean ± s.e.m. of at least n=20 individual cells, imaged on at least n=3 biologically independent experiments. The promoter used in these experiments was p4.

Versions of promoters p1 and p2 with an even stronger RE (promoters p10 and p11, respectively; Table S1) were fully activated by the low synTF levels present in the nucleus in the dark. Therefore, these constructs are not light-sensitive and have not been further analyzed.

### Weak coupling between TF binding and PIC assembly is the key feature that allows a promoter to distinguish dynamics

To gain a mechanistic understanding of the reasons why promoter p4 is the best at distinguishing synTF dynamics, while p1 cannot do it at all, we developed a compartmental mathematical model of gene expression based on ordinary differential equations (Figure 4A). We opted for a three-state promoter model, whereby the promoter can be inactive (synTF is unbound), active (synTF is bound to a certain fraction of the REs, the TATA binding protein (TBP) is bound to the TATA box and, consequently, the PIC can be assembled) and refractory (synTF is bound to the REs, but the PIC cannot be assembled). synTF can bind to all or a fraction of the REs present in the promoter. We model the DNA looping ––needed to bring the TF in close proximity to the TATA box for recruitment of the GTFs (Bartman et al., 2016; Petrascheck et al., 2005)–– by dividing the rate of PIC assembly by a factor, j_m_, that accounts for the distance between the last RE and the TATA box, given that the efficiency of DNA looping between two points decreases with increasing distance between them (Nolis et al., 2009; Ringrose et al., 1999). We finally assume that the maturation of the RNA and the fluorescent reporter protein occur at constant rates, while protein translation depends on mRNA concentration and ability of the ribosome to find the start codon (see Supplementary Text for the equations and a detailed description of the model).

**Figure 4.**
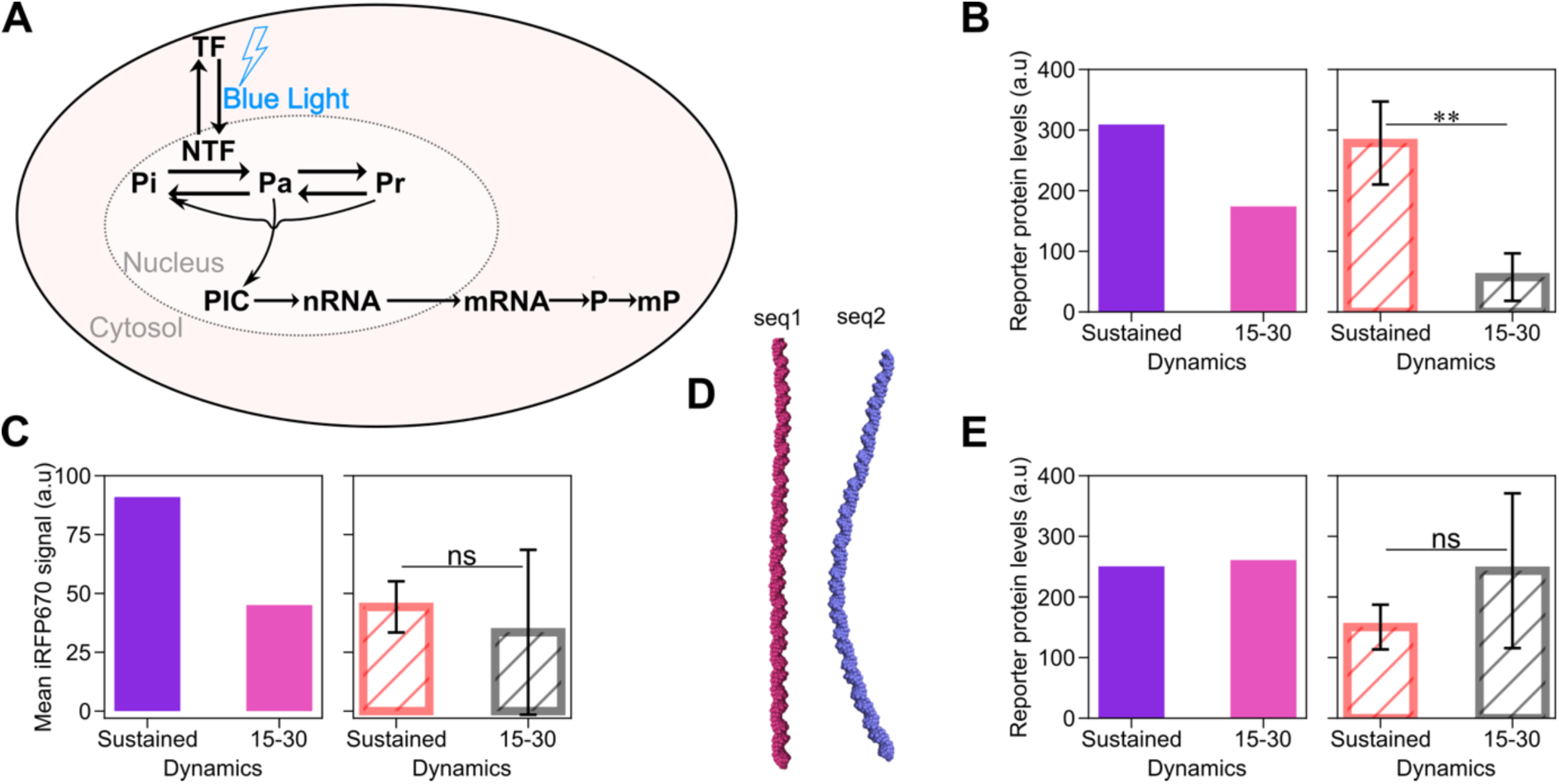
A mathematical model of gene expression mechanistically explains how a mammalian promoter can distinguish TF dynamics. (A) Schematic showing the reactions included in the mathematical model. The equations are explained in the Supplementary Text. (B,C,E) Model predictions (left panel; fitted (violet bar; for sustained dynamics) and simulated (pink bar; for both pulsatile dynamics)) and experimental data (right panel) of the mean reporter protein levels at the end of the experiment for the indicated synTF dynamics in combination with promoters p5 (B), p6 (C) and p7 (E). (D) Curvature of the indicated DNA sequences modeled using the cgDNAweb+ web server (De Bruin and Maddocks, 2018). (B,C,E) Data represent mean ± s.e.m. of at least n=20 individual cells, imaged on at least n=3 biologically independent experiments. P-values were calculated with the Welch’s t-test. ns, non signficant (P > 0.05). **, P-value = 0.00369.

We assigned physiologically reasonable values to some parameters based on the literature (Hahn et al., 1989; Zhang et al., 2010) and further used the data obtained with the sustained dynamics with promoter p1 to fit the remaining parameters (Figures 2A-2C, left panels). We then used it to simulate synTF dynamics, and mean reporter nascent RNA and protein levels over time for the two pulsatile dynamics with promoter p1. We found a good agreement between the simulations and the experimental data (Figures 2A-2C, middle and right panels).

Having now a mathematical model able to describe the behavior of promoter p1, we used it to understand why it gives rise to higher reporter protein levels at the end of the experiment, despite being insensitive to TF dynamics, as seen by the RNA data. We hypothesized that the reason for this behavior lied in the potential higher *effective* cumulative synTF levels of the pulsatile experiments, which run for a longer period of time than the experiment for sustained dynamics. Using the model, we determined the threshold for synTF concentration above which the reporter protein levels are above the half maximal value (Figure S2F) and used this value to quantify the effective cumulative levels for synTF for sustained and 15-30 pulses (Figure S2G). We found that, indeed, they are higher for the pulsatile dynamics. After normalizing the experimental mean reporter protein levels at the end of the experiments against the effective cumulative synTF levels found *in silico*, the difference among the dynamics disappears (Figure S2H).

The model successfully predicts the behavior of promoters p2-p4, when the promoter-specific parameters are updated using the sustained dynamics data (Figures S2A-2E, and Figures 3A-3C, left panels). Importantly, the models describing the four promoters are identical in the set of reactions, the only difference being the values of the promoter-specific parameters, which require a fitting step for each promoter.

Because the model of gene expression we developed captures all experimental data, we conclude that the processes we included are sufficient to explain how a mammalian promoter distinguishes TF dynamics. Our model offers the following mechanistic explanation: if the coupling between TF binding and PIC assembly is very efficient, any TF binding event, even a brief one, will be sufficient to recruit the GTFs, assemble the PIC and initiate transcription. The coupling is efficient, for instance, if the TF binds with high affinity to the RE, and/or the TBP binds with high affinity to the TATA box. A promoter, such as p4, with weak RE and TATA box, has inefficient coupling and, therefore, is less activated by pulsatile dynamics, characterized by a shorter residence time of the TF in the nucleus, and, consequently, at the promoter.

To demonstrate that the efficiency of coupling is the key feature that renders a mammalian promoter sensitive to TF dynamics, we used our mathematical model to predict whether we could turn promoter p1 from insensitive to sensitive to TF dynamics by extending the distance between the last RE and the TATA box (promoter p5). Indeed, as previously mentioned, the longer the distance, the less efficient the DNA looping, and, consequently, the coupling between TF binding and GTFs recruitment. After fitting the promoter-specific parameters using the data obtained with the sustained dynamics, we used the model to predict the behavior of promoter p5 under the pulses. The simulations indicate that promoter p5 is activated twice as much by sustained dynamics than low-frequency pulses (Figure 4B, left panel). We constructed promoter p5 by inserting a random DNA sequence of 147 bp (seq1) between the last RE and the TATA box in promoter p1 (λ=196 bp; Table S1 and Figure 1J). Despite producing overall much less reporter protein than promoter p1 under sustained dynamics (compare Figures 2E and 4B, right panel), promoter p5 barely responded to the 15-30 pulses (Figure 4B, right panel), implying that decreasing the coupling efficiency was sufficient to render promoter p1 sensitive to dynamics. When *in silico* introducing the same distance between the RE and the TATA box into promoter p2 (creating promoter p6), we found a similar trend, with sustained activation leading to higher protein production than pulses (Figure 4C, left panel). To validate the model predictions, we constructed promoter p6 by inserting the same random sequence seq1 into promoter p2 (Table S1). However, promoter p6 was too weak to be analyzed (Figure 4C, right panel).

Sequence length is, however, not the only important parameter for DNA looping. Reasoning that DNA looping efficiency could be improved by changing the DNA sequence keeping the same distance between the RE and the TATA box, we cloned promoter p7 using a sequence which was predicted to be prone to looping (Lowary and Widom, 1998) (seq2; Figure 4D). The model predicted that promoter p7 produces more protein than promoter p6 and that it is insensitive to dynamics (Figure 4E, left panel), which we experimentally confirmed (Figure 4E, right panel). This suggests that, as far as the coupling between TF binding and GTFs recruitment is efficient, even promoters with overall low activity can respond well to low-frequency pulses.

### Inefficient translation initiation strengthens the ability of a promoter to decode TF dynamics

As mentioned above, promoters p2 and p3 ––that contain four REs–– distinguish synTF dynamics, but this is visible only at the level of the RNA (Figures 2G, 2H, 2J and 2K). One explanation for this behavior is that, when the RNA is efficiently translated into protein, the difference at the RNA level gets lost at the protein level, when the RNA levels are high. This compensation does not occur anymore when the RNA levels are much lower: promoter p8, which is a version of promoter p2 with two instead of four REs that produces less RNA (reporter nascent RNA could not be detected with our visualization method), is able to sense dynamics also at the protein level (Figure S4). To test the hypothesis that translation efficiency might play a role in transmitting the information encoded in TF dynamics, we built a small library of reporter constructs, whereby the promoter is fixed, but the 5’UTR varies (Figure 1D and Table S1). In particular, we tested the role played by mRNA scanning by the ribosome to locate the start codon, not other processes involved in mRNA translation. We took promoter p2 and, as the first step, decreased the distance between the TATA box and the start codon ATG, creating construct 5UTR1 (Figure 5A). This should eliminate any secondary RNA structures that may affect protein translation (Haimov et al., 2015) and should increase the efficiency of the translation initiation complex to locate the start codon (Araujo et al., 2012; Hinnebusch, 2011, 2014; Leppek et al., 2018; Pestova and Kolupaeva, 2002). The second construct, 5UTR2, is similar to 5UTR1, with the only difference that we exchanged the G at position -6 relative to the start codon to A, which should reduce translation efficiency (De Angioletti et al., 2004; Mohan et al., 2014) (Figure 5A). Finally, we created a construct, 5UTR3, lacking the Kozak sequence (Figure 5A). We transfected these constructs in HEK293 cells, imposed the two light regimes for sustained and low-frequency pulsatile dynamics, and measured synTF nuclear concentration and mean reporter protein levels over time (Figures S5 and 5B). The first observation we made is that, indeed, decreasing the length of the 5’UTR has a positive effect on reporter gene expression, since construct 5UTR1 leads to twice as high mean reporter protein levels than the original construct with promoter p2 with the longer 5’UTR under both, sustained dynamics and 15-30 pulses (compare Figures 2H and 5C, left panel). Construct 5UTR2 gave rise to ~1.2 times lower mean reporter protein levels under both conditions (Figure 5C). Neither construct showed sensitivity to synTF dynamics. Construct 5UTR3, which is characterized by the lowest translation initiation efficiency due to complete lack of the Kozak consensus sequence, enabled distinguishing TF dynamics (Figure 5D).

**Figure 5.**
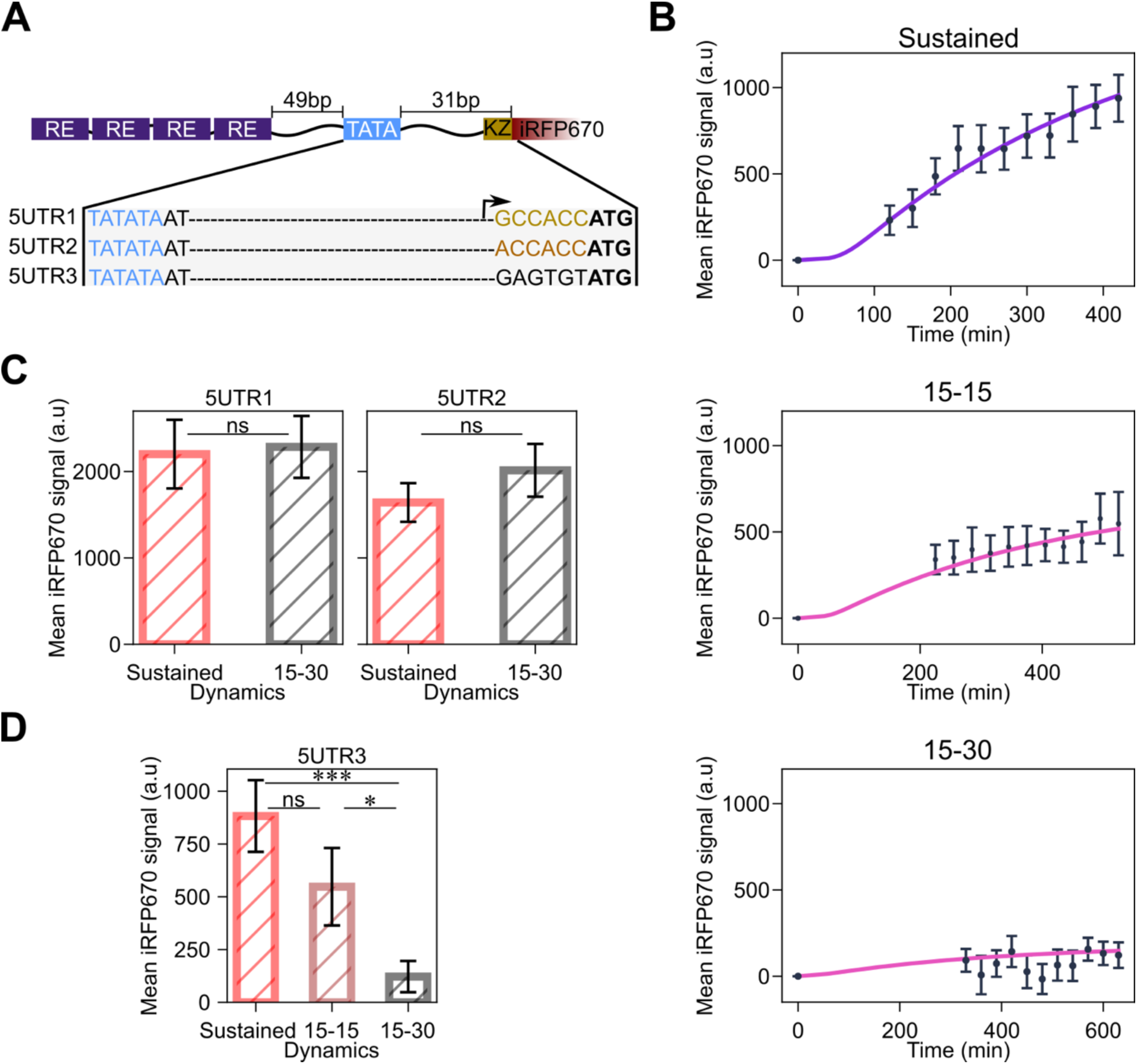
Decreasing translation initiation efficiency amplifies the ability of the promoter to sense TF dynamics. (A) Schematic showing the features of the constructs in the 5’UTR library. The Kozak sequence is shown in gold, the mutated Kozak sequence in orange, and the random sequence in black. The start codon is shown in bold, while the TATA box in blue. (B) Quantification of mean reporter protein levels over time for the indicated TF dynamics. Together with the experimental data (black dots), fitted (violet line; for sustained dynamics) and simulated (pink line; for both pulsatile dynamics) values are shown. The mathematical model used is shown in Figure 4 and the equations are described in the Supplememtary Text. (C,D) Quantification of reporter protein expression levels at the end of the experiment for the indicated synTF dynamics in combination with the indicated constructs. P-values were calculated with the Welch’s t-test. ns, non signficant (P > 0.05). *, P-value = 0.0419;.***, P-value = 0.0002. (B-C) Data represent mean ± s.e.m. of at least n=20 individual cells, imaged on at least n=3 biologically independent experiments.

Our mathematical model could correctly predict the behavior of the constructs in this library under the two pulsatile dynamics, after updating the translation parameters using the data from the sustained dynamics (Figures 5B and S5).

### Promoters activated by pulsatile but not sustained dynamics can theoretically exist

From our data and previous studies (Hansen and O’Shea, 2013; Li et al., 2018; Wilson et al., 2017), we can conclude that, whenever a promoter is activated by pulsatile dynamics, it is also activated by a sustained TF signal. In other words, from the building blocks that make a promoter (mostly the RE and the TATA box), it is not possible to obtain a variant that specifically filters sustained signals out, while responding to pulsatile ones. Wilson and colleagues showed, for the Ras/Erk pathway, that negative feedback can promote a band-pass filtering behavior allowing target genes to be most efficiently activated by Erk pulses of specific frequencies (Wilson et al., 2017). We asked ourselves if there could exist an alternative mechanism, involving no other molecule than the TF itself, which could allow a promoter to be better activated by pulsatile than sustained TF dynamics.

Recently, phase separation around genomic loci has been shown to play a regulatory role in gene expression (Cai et al., 2019; Sen et al., 2020; Shrinivas et al., 2019). These studies relate condensate formation with enhanced activity of the transcriptional activators that localize in them. Theoretically, however, formation of condensates could have an inhibitory function, as well: too high local TF concentrations could lead to a strong refractory response of the promoter, which could eventually enter into an inactive state. As a reminder, the refractory state is defined as that state of the promoter for which the PIC cannot be assembled despite the TF being bound at the RE(s), due to lack of GTFs available locally at the promoter to nucleate the PIC. We sought to explore *in silico* whether this mechanism could make a promoter respond better to pulsatile than sustained TF dynamics. We modified the original mathematical model to include a fourth promoter state: inactive. We assume that the inactive state is reachable from the refractory state and that, from this inactive state, the promoter can go back to being in the unbound state (Figure 6A). The rate at which the promoter switches from the inactive to the unbound state is a nonlinear inverse hill function of synTF concentration multiplied by the parameter D_in_, which is a scaling factor that depends on how fast the TF dissociates from the promoter. With this modified model we scanned the values for D_in_ and the interval between pulses that would lead to higher reporter protein levels at the end of the experiment (630 min) for pulses than sustained synTF signal (time between pulses = 0). We found several combinations of D_in_ and pulse frequencies that would lead to higher reporter protein levels in this case (Figure 6B). We then took one such combination and simulated two dynamics for synTF: sustained and pulses of the selected frequency specified by the arrow in Figure 6B (55 min light activation followed by 25 min dark phase; D_in_= 0.0031). Importantly, also in these simulations, we kept synTF cumulative levels constant, as done in the experiments. Therefore, the activation of synTF in the case of the pulsatile dynamics goes on for a longer time than for the sustained dynamics. We calculated the simulated mean nascent RNA levels over time for both dynamics. The model predicts that, while transcription rapidly decreases and eventually ceases for the sustained synTF signal, transcription goes on for the pulsatile synTF signal until there is no nuclear synTF (Figure 6C). The predicted cumulative nascent RNA levels are, therefore, higher for the pulsatile than the sustained dynamics (Figure 6D). The simulations show that the sustained synTF signal would lead to much lower reporter protein levels than the pulses (Figure 6E). Taken together, the mathematical model indicates that it is possible for a promoter to be more efficiently activated by a pulsatile than a sustained TF signal, provided the TF inhibits PIC assembly when above a certain threshold.

**Figure 6.**
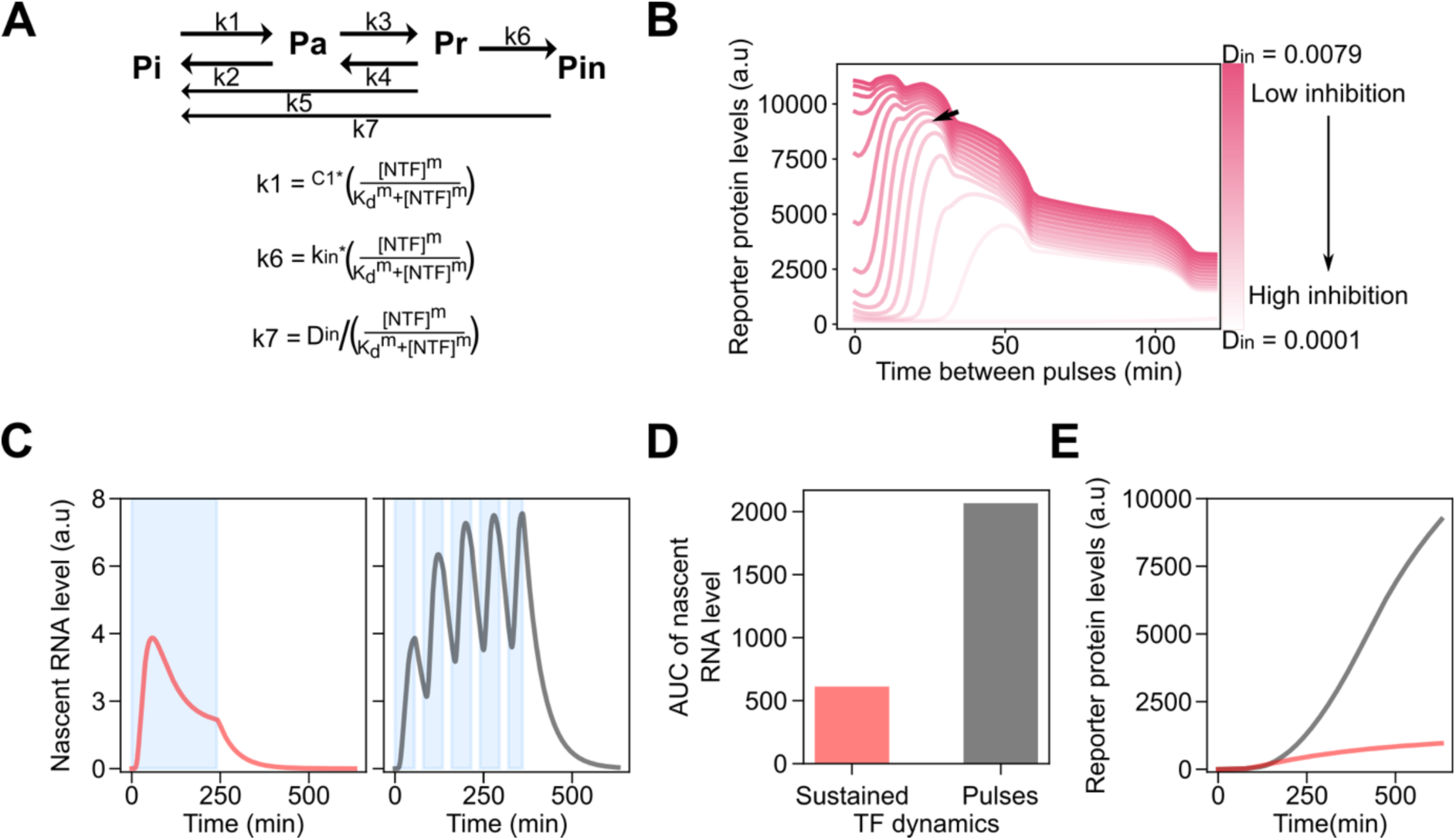
A promoter that filters out sustained signal but is activated by pulses can theoretically exist. (A) Modified mathematical model that includes a fourth promoter state. [NTF], Concentration of synTF in the nucleus. m, Hill coefficient. (B) Simulated values of the reporter protein at the end of the experiment as a function of the interval between pulses for different values of the parameter D_in_. When the time between pulses is zero the signal is sustained. Black arrow, selection of parameters used to simulate the system in (C-E). D_in_ = 0.0031. Time between pulses = 25 min. (C) Simulated nascent RNA levels for sustained (left panel) or pulsatile (right panel) dynamics using the model shown in (A) and the values for D_in_ and time between pulses selected in (B). Simulated area under the curve (AUC) for the nascent RNA for the indicated dynamics. (E) Simulated reporter protein levels over time for sustained (coral curve) or pulsatile (grey curve) TF dynamics.

## Discussion

In this study, we combined experimental and computational methods to decipher how mammalian promoters decode TF dynamics. We opted for a synthetic biology approach: we studied a synthetic TF (synTF) and synthetic promoters, all made of well-known and characterized parts. Being synTF orthogonal to the mammalian cells, it is not subject to endogenous regulation, which isolates our system from all other cellular processes and makes the data interpretation reliable. It additionally consented us to work quantitatively, knowing the affinities of the DBD and the TBP for the RE and the TATA box, respectively. It was interesting for us to see that a two-fold decrease in binding affinity between the TBP and the TATA box had a measurable effect. In our study, we call the TATA box bound by the TBP with K_D_ = 4 nM and the RE bound by synTF with K_D_ = 5.64 nM *weak*, despite these values being per se still in the nanomolar range.

To impose different dynamics to synTF we used an optogenetic tool, LINuS (Niopek et al., 2014), that allowed us to directly manipulate, with blue light, the nuclear concentration of synTF. We designed our experiments in a way that cells in all conditions would be exposed to the same total amount of light, permitting comparisons among the three investigated TF dynamics without differential interference of light.

To focus on TF dynamics, we decided to fix the cumulative TF levels, as previously done for p53 (Purvis et al., 2012). Some promoters, like promoter p1 in our library, are, indeed, sensitive to cumulative TF levels, rather than dynamics, and having different values for this parameter would confound the data interpretation. Although an alternative way to obtain similar cumulative levels across conditions would have been to fix the experimental time and change the amplitude of the signal in the three conditions, we opted against this choice, because some promoters are sensitive to TF amplitude, which would have precluded the possibility to isolate only effects due to dynamics. Since promoters p1 and p2 are insensitive to synTF amplitude (Figures 2F and 2I), we performed in this case also the experiment fixing the time and lowering the amplitude for the sustained dynamics to obtain similar cumulative synTF levels as for the pulses (Figure S2I). We found consistent results with those obtained with variable experimental time (Figures S2J-S2K), thus we concluded that our choice to fix cumulative levels by changing experimental time is valid.

In parallel to detecting the reporter protein, we established an RNA visualization method compatible with living cells to more directly monitor promoter activity. Nascent RNA measurements consented us to realize that promoters p2 and p3 distinguish dynamics (Figures 2G and 2J), although this is not seen at the protein level when the mRNA is efficiently translated (Figures 2H and 2K). The limitation of our setup ––that relies on mVenus to visualize nascent RNA–– is that we cannot take images in this channel during the dark phases of the pulsatile dynamics, because the illumination would lead to activation of LINuS within synTF and, consequently, to unwanted nuclear import and reporter gene expression. Despite this limitation, we expected the first visualization at the beginning of each blue light activation phase to be indicative of whether transcriptional shut down had occurred during the dark phase. Because nascent RNA levels after each dark incubation phase were similar to those obtained during blue light activation (Figures 2B, S2B and S2D, middle and right panels), we conclude that, at the population level, transcriptional shutdown during the phases in which synTF is in the cytoplasm does not occur. Looking at individual cells, we observed that some cells shut down, while others either don’t or do, but re-initiate RNA transcription during the dark phases (Movie S1). The appearance of RNA foci during the phases in which the synTF is not nuclear, and therefore unbound from the REs, could be explained by transcription re-initiation events that do not involve the TF (Joo et al., 2017; Yean and Gralla, 1999).

Nucleosome positioning has been previously suggested to play a critical role in the ability of a promoter to decode TF dynamics (Hansen and O’Shea, 2013). Clearly, the presence of a nucleosome at the promoter renders the coupling between TF binding events and PIC assembly inefficient, thus it is in line with our explanation of what makes a promoter sense TF dynamics. Nonetheless, our data indicate that there are other ways to achieve this, independent of the chromatin status.

We found particularly interesting the observation that lowering the translation initiation efficiency amplifies the ability of promoter p2 to sense dynamics. This is explained considering that mRNA translation and degradation compete with one another, with high ribosome occupancy protecting mRNA from degradation (Jia et al., 2020). If the mRNA is inefficiently translated, higher mRNA concentrations will be required to improve the chances of successfully initiating translation on high enough a fraction of mRNA molecules before they are degraded.

All the promoters in our library that respond to pulsatile synTF signals respond to the sustained signal, too. We highlight here that we did not explore cases of sustained signals of shorter total duration than pulses, or vice versa, but rather always compared different dynamics involving the same cumulative TF levels. Natural signaling pathways, however, do exist for which certain target genes are more efficiently activated by pulsatile than sustained signal. For the Ras/Erk pathway, for instance, it was shown that immediate-early genes (IEGs) respond better to Erk pulses of intermediate frequency than to sustained Erk (Wilson et al., 2017). The mechanism that allows IEGs to filter out a sustained Erk signal involves negative feedback enacted by Erk negative regulators. Our synthetic gene network does not comprise negative feedback; therefore, in our setup, promoters that are activated by pulses are also necessarily activated by sustained synTF signal. Using our mathematical model, we theoretically explored the possibility that sustained TF signals be filtered out via a mechanism not involving a protein other than the TF itself. We envisioned that this could happen if synTF residing in the nucleus for long enough would repress, instead of activate, the promoter. The model indicates that this mechanism would indeed allow accumulating higher reporter protein levels for pulsatile than sustained synTF dynamics (Figure 6). Although this mechanism has not been proved to occur in natural systems yet, it could be implemented using synthetic biology and protein engineering approaches.

In summary, here we provided a mechanistic understanding of the features that render mammalian promoters sensitive to TF dynamics. Our results will help interpret how natural signaling pathways are activated by different dynamic inputs as well as pave the way to construct synthetic circuits that respond differently to a single input molecule depending on its dynamics.

## Materials and Methods

### Bacterial strains for molecular cloning

Chemically competent *E. coli* TOP10 or DH5α cells were used for the transformation of circular plasmid DNA. For plasmid amplification, kanamycin or ampicillin was used as a selection agent at a final concentration of 50 µg ml^−1^ or 100 µg ml^−1^, respectively. All bacterial cells were incubated in lysogeny broth medium (LB) and on LB agar plates containing the appropriate antibiotic.

### Cell lines and maintenance

HEK293 cells were maintained at 37°C and 5% CO_2_ in phenol red-free Dulbecco’s Modified Eagle Medium (Gibco, Thermo Fisher Scientific) supplemented with 10% fetal calf serum (Sigma-Aldrich (MilliporSigma)), 2 mM L-glutamine (Gibco, Thermo Fisher Scientific), 100 U ml^-1^ penicillin and 10 mg ml^-1^ streptomycin (Gibco, Thermo Fisher Scientific).

### Molecular cloning

#### PCR for molecular cloning

Single-stranded primer deoxyribonucleotides with a final concentration of 100 μM were ordered from Sigma Aldrich or Eurofins Genomics. PCR reactions with plasmid and genomic DNA templates were performed using the Phusion High-Fidelity 2× Master Mix or Q5 High-Fidelity 2× Master Mix (New England Biolabs) according to the manufacturer’s protocol. Samples were purified by DNA agarose gel electrophoresis followed by gel extraction using QIAquick Gel Extraction Kit (Qiagen).

#### Molecular cloning using Gibson assembly

Plasmid pDN98 (Niopek et al., 2014) was used as parental plasmid for the construction of the synthetic TF (synTF). The LexA dimerization domain was amplified from genomic DNA extracted from *E. coli* (TOP10 strain) and inserted between the LexA DNA-binding domain (DBD) and the VP48 transactivation domain (TAD) in pDN98 to generate plasmid pEA00. The iRFP670 coding sequence was PCR-amplified from plasmid pNLS-iRFP670 (Shcherbakova and Verkhusha, 2013) (gift from Vladislav Verkhusha (Addgene plasmid #45466) with a primer containing the CAAX motif and cloned in place of the firefly luciferase gene into plasmid pDN100 (Niopek et al., 2014). The DNA sequence encompassing the synthetic promoter, the reporter gene and the bovine growth hormone (BGH) terminator was then PCR-amplified from this modified pDN100 plasmid and cloned into pEA00 upstream of the CMV promoter (synTF promoter) in a tandem orientation, giving rise to plasmid pEA01 (synPlasmid1). Insertion of the reporter construct in a convergent orientation downstream of the SV40 terminator led to the construction pEA02 (synPlasmid2). Reversing the orientation of the synTF construct in pEA01 gave rise to pEA03 (synPlasmid3). All other reporter library constructs were generated by modifying an element in the promoter of the synthetic reporter in pEA01.

The MS2coat protein (MCP) was amplified from plasmid ubc-nls-ha-MCP-VenusN-nls-ha-PCP-VenusC (Wu et al., 2014) (gift from Robert Singer; Addgene plasmid #52985). The IRES-SV40/NLS-MCP gene sequence together with full-length mVenus gene amplified from pTriEx-NTOM20-mVenus-Zdk2 (Wang et al., 2016) (gift from Klaus Hahn; Addgene plasmid #81011) was inserted after the stop codon of synTF in pEA01. The 12xMBS-PBS sequence was PCR-amplified from plasmid Pcr4-12xMBS-PBS (Wu et al., 2014) (gift from Robert Singer; Addgene plasmid #52984) and cloned after the stop codon of the reporter gene sequence. For better foci visualization, the BGH promoter was removed to allow for a longer 3’UTR, which permits the nascent RNA to be bound long enough for it to be visualized. The complete list of all plasmids is given in table S3. The list of all primers used in this study is given in table S4.

All plasmids were constructed using Gibson assembly (Gibson et al., 2009). Gibson assemblies were performed using 50 ng backbone DNA in a 5 µl reaction volume and a molar 1:1-3 backbone:insert ratio, using the NEBuilder HiFi DNA Assembly Master Mix (2×) (NEB) for 20–40 min at 50 °C. Agarose-gel purified DNA fragment concentrations were determined using a spectrophotometer (NanoDrop One, Thermo Fisher Scientific).

#### DNA agarose gel electrophoresis

Gels were prepared with 1% agarose (Agarose Standard, Carl Roth) in 0.5× TAE-buffer and 1:50,000 Ethidium Bromide (Roth), running for 20–30 min at 130 V. For analysis, 1 kb Plus DNA Ladder (NEB) was used. The samples were mixed with gel loading dye (purple, 6×) (NEB).

#### Bacterial transformation with plasmid DNA

Chemical transformation was performed by mixing 5 µl of Gibson reaction with 50 µl of chemically competent cells and incubating the mixture on ice for 30 min. Cells were then heat-shocked at 42 °C for 90 s, further incubated on ice for 5 min and finally mixed with 450 µl LB medium. Transformed cells were incubated on 37 °C shaker for 30-45 min before plating on agar plates containing the antibiotic. Plates were incubated overnight at 37 °C or for 48 h at room temperature.

#### Plasmid DNA purification and Sanger sequencing

Individual clones were picked from the agar plate and inoculated in 2-3 ml LB medium with kanamycin or ampicillin and incubated for about 6-8 h. Plasmid DNA was purified with the QIAprep Plasmid MiniSpin (QIAGEN) according to the manufacturer’s protocol. Plasmids were sent for Sanger sequencing (GATC-Biotech/Eurofins) and analyzed using APE (https://jorgensen.biology.utah.edu/wayned/ape/).

#### Plasmid transfection

Cells were transfected with the calcium phosphate transfection protocol. DNA amounts were kept constant in all of the experiments to yield reproducible complex formation and comparable results. A total amount of 500 ng of DNA was used to transfect cells seeded in Ibidi μ-dish 4-well dishes (Ibidi GmbH, Germany). pBlue-ScriptIIS/K was used as stuffer plasmid at a ratio of 1:200. Cells were plated 1 day before transfection (75,000 cells in 250 µl of culture medium per well of Ibidi μ-dish 4-well dishes (Ibidi GmbH, Germany)).

### Cellular imaging and optogenetic stimulation

Microscopy was performed always 24 hours post transfection. Cells were maintained at 37°C and 5% CO_2_ in a dark incubation chamber for the duration of the microscopy session. Images in the mCherry and iRFP670 channels were acquired in confocal modality on a Zeiss LSM 800 confocal microscope equipped with a motorized stage, a Plan-Apochromat 40x/1.4 numerical aperture oil immersion objective (Zeiss), a laser module containing 405, 488, 561 and 640 nm lasers and an electronically switchable illumination and detection module. Images in the mCherry channel were acquired with the following settings: 0.15% of 561 nm excitation laser using 37 μm pinhole aperture and 700 V gain. Images in the iRFP670 channel were acquired with 0.30% of 640 nm excitation laser using 41 μm pinhole aperture and 650 V gain. The confocal microscope was also equipped with a colibri light source consisting of 385, 425, 469, 511, 555 and 631 nm LEDs for widefield epifluorescence microscopy. Blue light activation was performed by exciting cells with 0.5% of 469 nm LED light in widefield microscopy mode using the 38 HE GFP filter set from Zeiss. The 0.5% light intensity, which corresponded to 6.44Wm^-2^ of light as measured with the LI-250A light sensor (LI-COR Biosciences), was achieved by filter out 75% of the 2% LED intensity. Automated cell focusing was done using mCherry as the reference channel. The sustained dynamics were generated by illuminating cells with blue light (GFP channel) for 125ms every 45s for 2h. The pulsatile dynamics were generated by illuminating the cells with the same illumination scheme used for sustained dynamics for 15 min, followed by a dark phase of either 15 (high-frequency pulses) or 30 (low-frequency pulses) min, and repeating this cycle 8x. For time-lapse tracking of synTF-mCherry localization, confocal mCherry images were taken every 5 min during activation and dark recovery phases. All image acquisitions were done using the ZenBlue software. mCherry and iRFP670 confocal images were taken prior to starting the optogenetic stimulation and post activation, every 30 min for 5h. Nascent RNA transcripts were visualized on the same microscope with an Axiocam503 camera and an 63x/1.4 Plan-Apochromat oil-immersion objective (Zeiss). Blue light activation of cells was performed with the same setup as above except that 0.95 % 469 nm LED light intensity was used to account for the change of objective. This LED intensity corresponded to 6.79 W m^-2^ of light. Images in the YFP channel (to image mVenus) were acquired using 5% of 511 nm LED light in widefield microscopy mode using the 46 HE YFP filter set from Zeiss in a Z-stack of 16-18 sections with 0.75 μm step size. Images were acquired every 5 min during the blue light activation phases. Since Z-stack imaging in the YFP channel was phototoxic to the cells when performed for a prolonged time, the pulsatile dynamics in the experiments for nascent RNA transcripts visualization was performed repeating the 15 min blue light activation six instead of eight times.

### Predicting DNA curvature and flexibility

Seq1 and seq2 were analysed using the cgDNAweb+ webserver (De Bruin and Maddocks, 2018) (https://cgdnaweb.epfl.ch/) with parameter set 4. The .pdb models were then aligned based on their coordinates using PyMol 2.4.1 (Delano, 2002).

### Modeling

Model simulations were performed in python v3.8.3.final.0 using the Anaconda v2020.07 distribution. Numerical simulations were performed using the odeint function in SciPy v1.5.0 scipy.integrate module, which is used as a wrapper for the LSODA ordinary differential equation solver for stiff or non-stiff systems from the FORTRAN library odepack. Initial conditions were set according to experimental data at time t=0 or were fitted from the sustained dynamics time-course data. The variables of interest were plotted using the matplotlib library.

### Quantification and statistical analysis

#### Image analysis

##### Automated quantification of mCherry and iRFP670 signals

The algorithm for automatic segmentation of the nucleus and cytosol in the absence of nuclear or plasma membrane markers has been described in detail in (Çiçek et al., 2020). Briefly, to segment the nucleus we apply the same base network as in (Çiçek et al., 2020) and use the off-the-shelf Mask R-CNN (He et al., 2017) trained with mCherry channel only. To augment the data and create nearly realistic input images, we also use the elastic deformations of U-Net (Ronneberger et al., 2015) to help improve the generalization capabilities of the network. To detect segmentation errors, we use both data uncertainty (aleatoric) and model uncertainty (epistemic). We model the former by learning the noise scale during training and computing the entropy of the class pseudo-probabilities for each pixel at test time as in (Kendall and Gal, 2017). For the later, we use the Winner-Takes-All (Ilg et al., 2018) method, which trains a single network with multiple heads and only updates the head with the best prediction every iteration. We choose this combination since it performs best in (Çiçek et al., 2020). To improve the output of Mask R-CNN, we compute the tracks as described in (Çiçek et al., 2020). We apply the suggested hyperparamters α = 0.7 and β = 0.85 to decide about frames that need to be updated. We consider a simple yet effective warping strategy by estimating the shift and scaling parameters computed between the not yet updated and neighbouring nuclei predictions. Likewise, we implicitly assume that the shape of the nuclei does not change over short time windows and only allow slight deformations to occur. Although flow-based methods tend to perform better according to (Çiçek et al., 2020), we do not use them to reduce the computational burden. To mitigate this slight drop in performance, we apply a sampling strategy before measuring the fluorescence. We report the average fluorescence of nucleus, cytosol and membrane per cell and frame. Instead of using the full prediction mask to compute the average, we sample a subset of pixels that have higher chances to belong to the corresponding structure. For nucleus and cytosol, segmentation errors occur mainly on the border. Therefore, we gradually erode the segmentation mask as long as it contains more than 2000 pixels. We then superpose the binary mask with mCherry channel and compute the average signal. Measuring the fluorescence for the membrane is very challenging since it is a very thin structure. Moreover, touching cells cause interference that amplify the signal. Thus, we use the iRFP670 channel and compute a skeleton. Notice that this skeleton might miss cells because of very low signal and might add artefacts in case of very high signal over surfaces. To avoid these errors, we rely on the cytosol masks and compute candidate pixels for the membrane by dilating the cytosol once. Then we remove intersections between candidate membranes of touching cells and pixels that are very close to the border of the image. Finally, we superpose the skeleton and the candidate membranes and compute the average based on the intersection. If there is no signal in the skeleton, we completely rely on the candidate membrane inferred from the cytosol.

##### Quantification of mVenus signal

The maximum projection of the Z-stacks was computed in ImageJ (Schneider et al., 2012) to bring all detected foci onto a single plane. To quantify nascent RNA, individual cells were first cropped and passed through a nascent RNA quantification pipeline as described in (Rullan et al., 2018). The first step in the pipeline was removal of the fluorescent background signal using a Gaussian filter. The filtered image was then subtracted from the original image to obtain an image which has the RNA foci features preserved without fluorescence background. A 2D Gaussian function was then fitted to the pixel intensity surface of each cell. The volume of the fitted function was used to represent the mean nascent RNA levels. The parameters of the 2D Gaussian function were limited to exclude fitting to background intensity fluctuations and large aggregates of the fluorescent proteins.

### Statistics and reproducibility

Statistics were calculated using the scipy.stats python module. No statistical method was used to predetermine sample size. The experiments were not randomized.

Dividing cells, cells that detached during the experiments and cells without well-defined nucleus at the start of the data acquisition were excluded from the analysis. Additionally, cells were stratified based on initial nuclear synTF levels and amplitude during blue light activation to allow for fair comparison of data from different dynamics and promoters. All experimental findings were reproduced in at least three independent experiments.

## Data availability

This study includes no data deposited in external repositories. Requests for materials should be directed to Barbara Di Ventura. All plasmids can be obtained under a Material Transfer Agreement.

## Acknowledgments

We thank Robert Grosse, Peter Walentek, Wolfgang Schamel and Johan Elf for critical reading of the manuscript, and members of the Di Ventura lab for useful discussions. We thank Vladislav Verkhusha for donating plasmid pNLS-iRFP670 (Addgene plasmid #45466), Robert Singer for donating plasmids Pcr4-12xMBS-PBS and Ubc-NLS-HA-MCP-mVenusN-NLS-HA-PCP-mVenusC (Addgene plasmids #52984 and #52985), and Klaus Hahn for donating plasmid pTriEx-NTOM20-mVenus-Zdk2(Wang et al., 2016) (Addgene plasmid #81011). This work was supported by the Federal Ministry of Education and Research – BMBF (grant no. 031L0079 to BDV), and by the Excellence Initiative of the German Federal and State Governments BIOSS (Centre for Biological Signalling Studies; EXC-294) and CIBSS (Center for Integrative Signalling Studies; EXC-2189).

## Author contributions

BDV conceived and supervised the study, and secured funding. EBA and BDV designed experiments and interpreted the data. EBA performed all experiments and mathematical modeling, wrote the first draft of the manuscript and prepared all figures. YM and ÖÇ developed the neural network for automated image analysis under the supervision of TB. BDV wrote the manuscript. All authors read and approved the manuscript.

