## Supplementary Material for "Requirements for mammalian promoters to decode transcription factor dynamics"

*for*


### Table of contents


**Supplementary Text**

**Tables S1-S4**

**Figs S1-S5**

**Supplementary Text**

***Mathematical model of gene expression in response to different synTF dynamics***

We model the process of gene expression using a deterministic, two-compartment model, with ten variables. The two compartments represent the cytosol and the nucleus, and the ten variables represent the following molecular species:

TF: cytosolic synTF
NTF: nuclear synTF
$P_i$: unbound promoter
$P_a$: active promoter
$P_r$: refractory promoter
PIC: pre-initiation complex
nRNA: nascent RNA
mRNA: matured mRNA
iP: immature protein
mP: mature protein

We opted for a deterministic, instead of a stochastic model of gene expression because our experimental setup is based on transient transfection of plasmid DNA, therefore, on average, there are more than two copies of the promoter of interest per cell; moreover, synTF is also expressed from the plasmid under the strong CMV promoter. We conclude that our species are present in medium copy and we can follow the average concentration of the species using ordinary differential equations (ODEs).

For each variable, we wrote an ODE that describes the change over time of its concentration. The model consists of the following ten ordinary differential equations (ODEs) describing cytosolic synTF (1), nuclear synTF (2), unbound promoter (3), active promoter (4), refractory promoter (5), PIC formation or stabilization (6), nascent RNA production (7), RNA maturation (8), mRNA translation to immature protein (9), protein maturation (10), plus one equation that describes DNA looping efficiency (11).

The first two equations describe the change over time in the concentration of synTF in the cytoplasm and the nucleus:

$$\frac{dTF}{dt} = K_{TF} + darkrev(t) - lightAct(t) - d_{TF} * [TF] \quad (1)$$

$$\frac{dNTF}{dt} = lightAct(t) - darkrev(t) - d_{NTF} * [NTF] \quad (2)$$

$K_{TF}$, $d_{TF}$ and $d_{NTF}$ are constants that describes the de novo production rate and degradation rates of synTF in the cytosol and nucleus, respectively. Production occurs only in the cytoplasm, while degradation occurs in both compartments.

darkrev(t) and lightAct(t) are two functions that describe the export and import of synTF from/into the nucleus, respectively.

The lightAct(t) function is dependent on time (t), duration of activating blue light (tOn), duration of recovery phase in the dark (tOff), rates of synTF nuclear import during light activation (iOn) and dark recovery phase (iOff). We added this latter nuclear import in the dark to take into account the fact that the LOV domain is in equilibrium between the caged, folded state, in which the NLS is shielded from the import machinery, and the uncaged, unfolded state that permits NLS recognition (*16*). The duration of the light activation (lightdur) defines the duration of synTF dynamics in minutes. The function lightAct(t) is defined in python as shown below and returns synTF import rate (imp):

```
def lightAct(t, tOn, tOff, iOn, iOff):
    if (t <= lightdur):
        if (t%(tOn + tOff) < tOn):
            imp = iOn
        elif (t%(tOn + tOff) >= tOn):
            imp = iOff
    else:
        imp = iOff
    return(imp )
```

The darkrev(t) function is dependent on time (t), duration of activating blue light on (tOn), duration of recovery phase in the dark (tOff), rates of synTF export out of the nucleus during light activation (rOn) and dark (rOff). The function darkrev(t) is defined in python as shown below and returns synNTF export rate (exp):

```
def darkrev(t, tOn, tOff, rOn, rOff):
    if (t <= lightdur):
        if (t%(tOn + tOff) < tOn):
            exp = rOn
        elif (t%(tOn + tOff) >= tOn):
            exp = rOff
    else:
        exp = rOff
    return(exp)
```

The next three equations describe the change over time in the promoter state. The promoter can assume three states: unbound ($P_i$), active ($P_a$) and refractory ($P_r$). Only in the active state the promoter is able to trigger transcription pre-initiation complex (PIC) assembly (described by equation 6). A three-state promoter model that includes a refractory state was necessary to account for the refractory behaviour seen in the nascent RNA data of some promoters. All promoter states are reversible.

Equation 3 describes the species $P_i$. $P_i$ becomes $P_a$ when bound by NTF. NTF binding to $P_i$ is modelled as cooperative using a Hill function of NTF concentration multiplied by the constant $K_{on}$, where m is the Hill coefficient and $kD_1$ is the affinity of synTF for the RE. The change over time of $P_i$ concentration also depends of the rate at which $P_a$ and $P_r$ revert back to the unbound state $P_i$ ($K_{off}$ and $d1_{rf}$, respectively):

$$\frac{dP_i}{dt} = K_{off} * P_a + d1_{rf} * P_r - K_{on} * \left(\frac{NTF^m}{{kD_1}^m + NTF^m}\right) * P_i \quad (3)$$

Equation 4 describes the species $P_a$. $P_a$ is formed when $P_i$ is bound by synTF. As mentioned above, NTF binding to $P_i$ is modelled to be cooperative using a Hill function of NTF concentration multiplied by the constant $K_{on}$, where m is the Hill coefficient and $kD_1$ is the affinity of synTF for the RE. $P_a$ is additionally gained back when $P_r$ spontaneously reverts back from refractory to active with rate $d2_{rf}$. $P_a$ can switch to either $P_i$ or $P_r$ with rates $K_{off}$ and $K_{rf}$, respectively:

$$\frac{dP_a}{dt} = K_{on} * \left(\frac{NTF^m}{{kD_1}^m + NTF^m}\right) * P_i + d2_{rf} * P_r - (K_{off} + K_{rf}) * P_a \quad (4)$$

Equation 5 describes the species $P_r$. Only species $P_a$ can become $P_r$ with rate $K_{rf}$, while $P_r$ can revert back to either $P_i$ or $P_a$ with rates $d1_{rf}$ and $d2_{rf}$, respectively:

$$\frac{dP_r}{dt} = K_{rf} * P_a - \left(d1_{rf} + d2_{rf}\right) * P_r \quad (5)$$

The sum of $P_i$, $P_a$ and $P_r$ was modelled as a constant throughout the simulation. For a promoter with 4 REs, for example, the total was 4 multiplied by a factor 2 – as adjustment for the values in the model.

Equation 6 describes the assembly of the PIC. This occurs when the promoter is in the active state $P_a$. The rate of PIC assembly depends on NTF binding to one or more REs, on the looping efficiency ($j_m^{-1}$) of the DNA sequence between the REs and the TATA box. We model this as a Hill function of NTF concentration, with Hill coefficient m and rate constant $K_{pic}$. The PIC can additionally disassemble with rate $d_{pic}$:

$$\frac{dPIC}{dt} = \frac{K_{pic}}{j_m} * \left(\frac{NTF^m}{{kD_1}^m + NTF^m}\right) * P_a - d_{pic} * PIC \quad (6)$$

Equation 7 describes RNA transcription of the target gene, that is, formation of nascent RNA (nRNA). We model this process as a Hill function of the PIC concentration, where n is the Hill coefficient and $kD_2$ is the dissociation constant of the PIC components from the core promoter, multiplied by the rate constant $K_{nrna}$. We use a Hill function to model the cooperative binding of the general transcription factors that form the PIC. Additionally, the nRNA is released from the DNA template at rate $d_{nRNA}$:

$$\frac{dnRNA}{dt} = K_{nrna} * \left(\frac{[PIC]^n}{{kD_2}^n + [PIC]^n}\right) - d_{nRNA} * nRNA \tag{7}$$

Equation 8 describes the formation of mature RNA (mRNA). This is a function of the nascent RNA that is processed and translocated to the cytosol as matured mRNA (mRNA) with rate $K_{mRNA}$. We also assume that the mRNA gets degraded at a constant rate $d_{mRNA}$:

$$\frac{dmRNA}{dt} = K_{mRNA} * nRNA - d_{mRNA} * mRNA \tag{8}$$

Equation 9 describes mRNA translation to an immature protein iP. We model this process as a Hill function of the mRNA concentration multiplied by the constant $K_{p1}$. We use a Hill function to model the fact that translation initiation is a cooperative process dependent on the affinity of ribosomal subunits for the start codon ($kD_3$). q is the Hill coefficient. iP becomes mature protein mP at rate Rp and is degraded at rate $d_p$:

$$\frac{diP}{dt} = K_{p1} * \left(\frac{[mRNA]^q}{{kD_3}^q + [mRNA]^q}\right) - \left(R_p + d_p\right) * iP \tag{9}$$

Finally, equation 10 describes the maturation of the reporter protein mP. mP gets degraded at rate $d_{mp}$:

$$\frac{dmP}{dt} = R_p * iP - d_{mP} * mP \tag{10}$$

The experimentally measured iRFP670 signal corresponds to the species mP.

Equation 11 describes how we calculate the factor $j_m$, and is based on previous work (*17*). L represents the distance between the TATA box and RE in bp, while P is the length of the DNA in nm. To calculate the length we use a previous estimate (*17*, *18*):

$$j_m = \left(\frac{1.25e^5}{P^3}\right) * \left(\frac{4*P}{L*10^4}\right)^{\frac{3}{2}} * e^{-\left(510*P^2\right)/\left(6.25*L^2+50*P^2\right)} \tag{11}$$

The ODEs were written in python v3.8.3.final.0 using the Anaconda v2020.07 distribution. Numerical simulations were performed using the odeint function in SciPy v1.5.0 scipy.integrate module, which is used as a wrapper for the LSODA ordinary differential equation solver for stiff or non-stiff systems from the FORTRAN library odepack.

Initial conditions were set according to experimental observations or were fitted. Variables of interest were plotted using matplotlib plotting library.

#### *Parameters*

The model entails a total of 21 parameters, if we consider the promoter-specific parameters. Parameters $kD_1$, m, $kD_2$, n and $R_p$ were fixed based on the RE, the TATA box and the maturation half-life of iRFP670, respectively. Parameters $K_{on}$, $K_{off}$, $K_{rf}$, $d1_{rf}$, $d2_{rf}$, $K_{pic}$, $d_{pic}$, $K_{nrna}$, and $d_{nRNA}$ were fitted using the sustained synTF dynamics nRNA data from promoter p1 using both adaptive memory programming for global optimization (ampgo) and basinhopping global optimization algorithms in the lmfit.minimize package in python.

While keeping q (Equation 10) at a value of 1, $K_{mRNA}$, $d_{mRNA}$, $K_{p1}$, $kd_3$, $d_p$, and $d_{mp}$ were fitted to the protein data obtained with the sustained synTF dynamics with promoter p1 using the same algorithms. The parameter set that successfully predicted the nascent RNA and protein data for the remaining pulsatile dynamics (high- and low-frequency) was used for the other promoters. For all other promoters, the only parameters that were allowed to change, beyond $kd_1$, m, $kd_2$ and n, were: $K_{on}$, $K_{off}$, $K_{rf}$, $d1_{rf}$ and $d2_{rf}$.

All other parameters were kept constant except when the number of REs or the 5’UTR changed. For promoters with 2 REs, $K_{pic}$, and $d_{pic}$ were allowed to change while all other parameters remained the same. We assume that a reduction in the number of REs affects PIC assembly and stabilization. In the instances of modified 5’UTR, only $K_{p1}$, $kD_3$ and q were allowed to change, since we reasoned that changing 5’UTR affects the translation efficiency of the mRNA, but not RNA transcription.

#### *Parameter fitting*

All parameter fittings were done with data from the sustained dynamics. Parameters were first fitted with the least square function (leastsq) from the lmfit.minimize python module. The resulting parameters were then used as the initial parameter guesses for global parameter fit using the adaptive memory programming for global optimization method (ampgo) or basinhopping algorithm implemented in lmfit.minimize python module. To improve on the global parameter values, the ampgo fitted output was used as initial guesses for another round of leastsq function fitting to find the local minima of the global parameters.

**Table S1** Complete list of reporter constructs tested in this study

| Promoter | #RE | LexA affinity for RE ($K_{D1}$) (nM) | λ (bp) | TBP affinity for TATA box $K_{D2}$ (nM) | δ (bp) | REF |
|---|---|---|---|---|---|---|
| **p1** | 4 | 1.61 | 49 | 2 | 69 | (*19*, *20*) |
| **p2** | 4 | 1.61 | 49 | 4 | 69 | (*19*, *20*) |
| **p3** | 4 | 5.64 | 49 | 2 | 69 | (*20*, *21*) |
| **p4** | 4 | 5.64 | 49 | 4 | 69 | (*20*, *21*) |
| **p5** | 4 | 1.61 | 196[a] | 2 | 69 | (*19*, *20*) |
| **p6** | 4 | 1.61 | 196[a] | 4 | 69 | (*19*, *20*) |
| **p7** | 4 | 1.61 | 196[b] | 4 | 69 | (*19*, *20*) |
| **p8** | 2 | 1.61 | 49 | 4 | 69 | (*19*, *20*) |
| **p9** | 2 | 1.61 | 49 | 2 | 69 | (*19*, *20*) |
| **p10** | 4 | 0.80 | 49 bp | 2 | 69 | (*20*, *21*) |
| **p11** | 4 | 0.80 | 49bp | 4 | 69 | (*20*, *21*) |
| **p12** | 2 | 0.80 | 49bp | 4 | 69 | (*20*, *21*) |
| **p13** | 4 | 1.61 | 343[c] | 4 | 69 | (*19*, *20*) |
| **p14** | 4 | 1.61 | 343[c] | 2 | 69 | (*19*, *20*) |
| **p15** | 4 | 0.80 | 343[c] | 2 | 31 | (*20*, *21*) |
| **p16** | 8 | 1.61 | 343[c] | 4 | 69 | (*19*, *20*) |
| **p17** | 4 | 1.61 | 343[d] | 4 | 69 | (*19*, *20*) |
| **p18** | 2 | 1.61 | 196[a] | 4 | 69 | (*19*, *20*) |
| **p19[e]** | 4 | 1.61 | 49 | N/A | 69 | (*19*, *20*) |
| **p20[f]** | 4 | 1.61 | 49 | N/A | 31 | (*19*, *20*) |
| **p21[g]** | 4 | 1.61 | 49 | N/A | 69 | (*19*, *20*) |
| **5UTR1** | 4 | 1.61 | 49 | 4 | 31[h] | (*19*, *20*) |
| **5UTR2** | 4 | 1.61 | 49 | 4 | 31[i] | (*19*, *20*) |
| **5UTR3** | 4 | 1.61 | 49 | 4 | 31[j] | (*19*, *20*) |
| **5UTR4** | 4 | 1.61 | 49 | 4 | 31[k] | (*19*, *20*) |
| **5UTR5** | 4 | 1.61 | 49 | 4 | 31[l] | (*19*, *20*) |
| **5UTR6** | 4 | 1.61 | 49 | 2 | 31[m] | (*19*, *20*) |
| **5UTR7** | 4 | 5.64 | 49 | 4 | 31[n] | (*20*, *21*) |
| **5UTR8** | 4 | 1.61 | 49 | N/A | 31[o] | (*19*, *20*) |

[a] Insertion of seq1 between REs and TATA box. [b] Insertion of seq2 between REs and TATA box. [c] Insertion of 2x seq1 between REs and TATA box. [d] Insertion of 2x seq1 flanked by the CTCF binding sequence on both 5' and 3' ends. [e] Promoter with initiator sequence in place of TATA box combined by downstream promoter element. [f] Promoter with 2x initiator sequence (Inr) in place of the TATA box. [g] Promoter with 2x initiator sequence in place of the TATA-box. [h] Promoter p2 combined with optimal Kozak sequence. [i] Promoter p2 combined with suboptimal Kozak sequence. [j] Promoter p2 combined with a random sequence in place of the Kozak sequence. [k] Promoter p2 with 62% GC content between

the TATA box and the start codon without Kozak sequence. [l] Promoter p2 with 55% GC content between the TATA box and the start codon combined with the suboptimal Kozak sequence. [m] Promoter p1 with 59% GC content between the TATA box and the start codon, and optimal Kozak sequence. [n] Promoter p4 with 59% GC content between the TATA box and the start codon, and optimal Kozak sequence. [o] Promoter p1 with the start codon, and optimal Kozak sequence without the TATA box.

**Table S2** Model parameters

| *Parameter* | ***p1*** | ***p2*** | ***p3*** | ***p4*** | ***p5*** | ***p6*** | ***p7*** | ***p8*** |
|---|---|---|---|---|---|---|---|---|
| *kD1* | 161.1805 | 161.1805 | 564.0000 | 564.0000 | 161.1805 | 161.1805 | 161.1805 | 161.1805 |
| *m* | 2.4588 | 2.4588 | 3.2487 | 3.2487 | 2.4588 | 2.4588 | 2.4588 | 2.4588 |
| *kD2* | 200.0000 | 400.0000 | 200.0000 | 400.0000 | 200.0000 | 400.0000 | 400.0000 | 400.0000 |
| *n* | 1.0000 | 3.2224 | 1.0000 | 3.2224 | 4.4668 | 3.1351 | 2.0000 | 3.2224 |
| *Kon* | 0.3936 | 0.3936 | 0.3149 | 0.3921 | 0.0080 | 0.0080 | 0.0080 | 0.0192 |
| *Koff* | 0.0194 | 0.0176 | 0.0200 | 0.0199 | 0.0011 | 0.0010 | 0.0020 | 0.0020 |
| *Krf* | 0.7060 | 0.1274 | 0.1980 | 0.0532 | 0.1274 | 0.1274 | 0.1274 | 0.1274 |
| *d1rf* | 2.36E-02 | 3.88E-03 | 1.99E-03 | 2.99E-08 | 4.62E-07 | 1.01E-04 | 1.00E-04 | 1.00E-04 |
| *d2rf* | 4.67E-03 | 1.04E-04 | 3.71E-05 | 5.68E-09 | 2.53E-05 | 2.79E-05 | 1.00E-04 | 1.00E-04 |
| *Kpic* | 1.07E-04 | 1.07E-04 | 1.07E-04 | 1.07E-04 | 1.07E-04 | 1.07E-04 | 1.07E-04 | 1.07E-04 |
| *dpic* | 0.2012 | 0.2012 | 0.2012 | 0.2012 | 0.2012 | 0.2012 | 0.2012 | 0.2012 |
| *Knrna* | 0.1782 | 0.1782 | 0.1782 | 0.1782 | 0.1782 | 0.1782 | 0.1782 | 0.1782 |
| *dnRNA* | 0.0202 | 0.0202 | 0.0202 | 0.0202 | 0.0202 | 0.0202 | 0.0202 | 0.0202 |
| *KmRNA* | 0.5893 | 0.5893 | 0.5893 | 0.5893 | 0.5893 | 0.5893 | 0.5893 | 0.5893 |
| *dmRNA* | 0.0673 | 0.0673 | 0.0673 | 0.0673 | 0.0673 | 0.0673 | 0.0673 | 0.0673 |
| *Kp1* | 16.074 | 16.074 | 16.074 | 16.074 | 16.074 | 16.074 | 16.074 | 16.074 |
| *q* | 1.000 | 1.000 | 1.000 | 1.000 | 1.000 | 1.000 | 1.000 | 1.000 |
| *kD3* | 4.25E+01 | 4.25E+01 | 4.25E+01 | 4.25E+01 | 4.25E+01 | 4.25E+01 | 4.25E+01 | 4.25E+01 |
| *dp* | 3.95E-04 | 3.95E-04 | 3.95E-04 | 3.95E-04 | 3.95E-04 | 3.95E-04 | 3.95E-04 | 3.95E-04 |
| *dmp* | 7.37E-06 | 7.37E-06 | 7.37E-06 | 7.37E-06 | 7.37E-06 | 7.37E-06 | 7.37E-06 | 7.37E-06 |
| *Rp* | 2.31E-03 | 2.31E-03 | 2.31E-03 | 2.31E-03 | 2.31E-03 | 2.31E-03 | 2.31E-03 | 2.31E-03 |

**Table S2 cont.**

| *Parameter* | ***5UTR1*** | ***5UTR2*** | ***5UTR3*** |
|---|---|---|---|
| *kD1* | 161.1805 | 161.1805 | 161.1805 |
| *m* | 2.4588 | 2.4588 | 2.4588 |
| *kD2* | 400.0000 | 400.0000 | 400.0000 |
| *n* | 3.2224 | 3.2224 | 3.2224 |
| *Kon* | 0.3936 | 0.3936 | 0.3936 |
| *Koff* | 0.0176 | 0.0176 | 0.0176 |
| *Krf* | 0.1274 | 0.1274 | 0.1274 |
| *d1rf* | 3.88E-03 | 3.88E-03 | 3.88E-03 |
| *d2rf* | 1.04E-04 | 1.04E-04 | 1.04E-04 |
| *Kpic* | 1.07E-04 | 1.07E-04 | 1.07E-04 |
| *dpic* | 0.2012 | 0.2012 | 0.2012 |
| *Knrna* | 0.1782 | 0.1782 | 0.1782 |
| *dnRNA* | 0.0202 | 0.0202 | 0.0202 |
| *KmRNA* | 0.5893 | 0.5893 | 0.5893 |
| *dmRNA* | 0.0673 | 0.0673 | 0.0673 |

| | | | |
|---|---|---|---|
| *Kp1* | 98.451 | 71.880 | 98.451 |
| *q* | 2.596 | 2.596 | 9.116 |
| *kD3* | 4.25E+01 | 4.25E+01 | 4.25E+01 |
| *dp* | 3.95E-04 | 3.95E-04 | 3.95E-04 |
| *dmp* | 7.37E-06 | 7.37E-06 | 7.37E-06 |
| *Rp* | 2.31E-03 | 2.31E-03 | 2.31E-03 |

**Table S3** List of plasmids used in this study

| Name | Backbone | Insert | Promoter | Source |
|---|---|---|---|---|
| **pDN98** | pmCherry-N1 | LexA DNA binding /VP48 /IkBa NES/mCherry/LINuS (biNLS10) | CMV | (*1*) |
| **pDN100** | pFR-Luc | Firefly luciferase | 4x LexA operator based promoter | (*1*) |
| **pEA00** | pDN98 | Full length LexA | CMV | This study |
| **pEAXX** | pDN100 | iRFP670-CAAX | 4x LexA operator based promoter | This study |
| **pEA01** | pEA00 | 4x-LexA0/iRFP670-CAAX/BGH terminator | p2 | This study |
| **pEA02** | pEA00 | Reversed 4x-LexA0/iRFP670-CAAX/BGH terminator | p2 | This study |
| **pEA03** | pEA01 | Reversed CMV-full length LexA /VP48 /IkBa NES/mCherry/LINuS (biNLS10)/ SV40 terminator | p2 | This study |
| **pEA04** | pEA01 | Promoter p1 | - | This study |
| **pEA05** | pEA04 | Promoter p3 | - | This study |
| **pEA06** | pEA05 | Promoter p4 | - | This study |
| **pEA07** | pEA04 | Promoter p5 | - | This study |
| **pEA08** | pEA01 | Promoter p6 | - | This study |
| **pEA09** | pEA01 | Promoter p7 | - | This study |
| **pEA10** | pEA04 | Promoter p8 | - | This study |
| **pEA11** | pEA01 | Promoter p9 | - | This study |
| **pEA12** | pEA01 | Promoter p10 | - | This study |
| **pEA13** | pEA12 | Promoter p11 | - | This study |
| **pEA14** | pEA12 | Promoter p12 | - | This study |
| **pEA15** | pEA01 | Promoter p13 | - | This study |
| **pEA16** | pEA04 | Promoter p14 | - | This study |
| **pEA17** | pEA12 | Promoter p15 | - | This study |
| **pEA18** | pEA15 | Promoter p16 | - | This study |
| **pEA19** | pEA15 | Promoter p17 | - | This study |
| **pEA20** | pEA08 | Promoter p18 | - | This study |
| **pEA21** | pEA01 | Promoter p19 | - | This study |
| **pEA22** | pEA01 | Promoter p20 | - | This study |
| **pEA23** | pEA04 | Promoter p21 | - | This study |
| **pEA24** | pEA01 | Promoter p2 5UTR1 | - | This study |

| | | | | |
|---|---|---|---|---|
| **pEA25** | pEA01 | Promoter p2 5UTR2 | - | This study |
| **pEA26** | pEA01 | Promoter p2 5UTR3 | - | This study |
| **pEA27** | pEA01 | Promoter p2 5UTR4 | - | This study |
| **pEA28** | pEA01 | Promoter p2 5UTR5 | - | This study |
| **pEA29** | pEA04 | Promoter p2 5UTR6 | - | This study |
| **pEA30** | pEA06 | Promoter p2 5UTR7 | - | This study |
| **pEA31** | pEA01 | Promoter p2 5UTR8 | - | This study |
| **pEAm** | pEA01 | IRES-SV40/NLS-MCP | - | This study |
| **pEAm00** | pEAm | 12xMBS-PBS | - | This study |
| **pEAm01** | pEAm00 | Minus BGH terminator | - | This study |
| **pEAm02** | pEAm01 | Promoter p1 | - | This study |
| **pEAm03** | pEAm01 | Promoter p3 | - | This study |
| **pEAm04** | pEAm01 | Promoter p4 | | |
| **pEAm05** | pEAm01 | Promoter p5 | - | This study |
| **pEAm06** | pEAm01 | Promoter p6 | - | This study |
| **pEAm07** | pEAm01 | Promoter p7 | - | This study |
| **pEAm08** | pEAm01 | Promoter p8 | - | This study |
| **pEAm09** | pEAm01 | Promoter p9 | - | This study |
| **pEAm10** | pEAm01 | Promoter p12 | - | This study |

**Table S4** List of primers

| # | Primer sequence 5’-3’ |
|---|---|
| **1** | tcgtgtggctgccggtgaaccacttctggcgcaacagcat |
| **2** | gtcggccggcccgccgctttcgttaattaagctggttccgctaccacccagccagtcgccgttgcg |
| **3** | gaaagcggcgggccggcc |
| **4** | tggttcaccggcagccac |
| **5** | aaaagaagaaaaagaagtcaaagacaaagtgtgtaattatgtaggcggccgctcgagcatg |
| **6** | ggtggcgcctatttaccaac |
| **7** | gttggtaaataggcgccaccatggcgcgtaaggtcgatc |
| **8** | ttgtctttgacttctttttcttcttttaccccttatagcgttggtggtgggcggcgg |
| **9** | atagtaatcaattacggggtcattagttc |
| **10** | taataactaatgcatggcggtaatac |
| **11** | ccgccatgcattagttattacagacggatcgggagatc |
| **12** | accccgtaattgattactatgctggcaagtgtagcggtc |
| **13** | atccccgggtaccgagctcgaattccagcttggca |
| **14** | gctcggtacccggggatccccttttatagcgtctagagtctccgctcggactcg |
| **15** | tgatcagacatgtatattggactgtaaaaaaaaacagtggttatatgtacagactagactgtaaaaaaaaacagtggttatatgtacagactagactcgagtccgag |
| **16** | tccaatatacatgtctgatcactgtttttttttacagtctagtctgtacatataaccactgtttttttttacagtctagatgcggccgcgaa |
| **17** | ttaatacaacgaacggtgatgttgtcatagattcggcacatttcccttgtaggtgtgaaatcacttagcttcgcgccgaagtcttatgagtccgagcggagactc |
| **18** | tgacaacatcaccgttcgttgtattaatcgatggtggtagcggctgcatattgtcagatgaaggagcgacacccggggaggagtccagagtctagtctgtacata |
| **19** | cgtacgcgctgtcccccgcgttttaaccgccaaggggattactccctagtctccaggcacgtgtcagatatatacatcctgatgagtccgagcggagactcta |
| **20** | cgcgggggacagcgcgtacgtgcgtttaagcggtgctagagctgtctacgaccaattgagcggcctgcagaccgggattctccagagtctagtctgtacatata |
| **21** | aattcgcggccgcatctagactgtatataaaaccagtgatcagacatgtatattggactgtatataaaaccagtggttatatgtacagactagactcgagtccgagcg |
| **22** | tctagatgcggccgcgaattcggta |
| **23** | tatatacagtgatcagacatgtatattggactgtatatatatacagtggttatatgtacagactagactgtatatatatacagtggttatatgtacagactagactcgagtccg |
| **24** | tccaatatacatgtctgatcactgtatatatatacagtctagtctgtacatataaccactgtatatatatacagtctagatgcggccgcgaattcgg |
| **25** | gcgtagctgcgcataagcaaatgacaattaaccactgtgtactcgttataacatctggcagttaaagtcgggagaataggagccgagtccgagcggagactc |
| **26** | ttgcttatgcgcagctacgccatcgcgaggccggtccggcgggcgaagcatataaaagaagctcgtcacatccacatagttgtataagacttcggcgcgaa |
| **27** | tgatcagacatgtatattggactgtatataaaaccagtggttatatgtacagactagactgtatataaaaccagtgattcgcggccgcatctagactgt |
| **28** | tccaatatacatgtctgatcactggttttatatacagtctagtctgtacatataaccactggttttatatacagtcggtacccggtcacagcttgtctg |
| **29** | gactcttactccctagtcttggatccccgggtaccg |
| **30** | aagactagggagtaagagtctccgctcggactc |
| **31** | gactcttactccctagtctttccggtactgttggtaaatagg |
| **32** | atggcgcgtaaggtcgatctcacctcctgcgatcgcgagccg |
| **33** | atcgaccttacgcgccatggtggcaagagttgcttcgtgcatagccgattatataccctc |
| **34** | atcgaccttacgcgccatggtggttagagttgcttcgtgcatagccgattatataccctc |
| **35** | atcgaccttacgcgccatacactcaagagttgcttcgtgcatagccgattatataccctc |
| **36** | atggatccccgggtaccgagctcgaattccaatggcgcgtaaggtcgatc |
| **37** | tggaattcgagctcggtacccggggatccattatataccctctagagtctccgctc |
| **38** | atcgaccttacgcgccatggtggttcgagttgattcgtgcatagccgattatataccctctagagtctccgctcggactcg |
| **39** | tggcgcctatttaccaacagtaccgcccttttatagcgtctagagtctccgctcggactc |
| **40** | ggtggcgcctatttaccaacagtaccggaatcgcgcgccctctagagtctccgc |
| **41** | TCGCTCGCTCCAGTATTCCAGGGTTCATCAGAGCATGCATCTAGAGGGCC |

| | |
|---|---|
| **42** | TGCTTTCTTGGCAATAAGTACCGTAGGATCACTAGTACTTCCACCTGAACCTCCctacataattacacactttgtctttgac |
| **43** | acctaaatgctagagctcgctgatcagcctatagtaatcaattacggggtc |
| **44** | AGGCTGATCAGCGAGCTC |
| **45** | agaagaaaaagctggactagATCGATGGATCCCTCCCCCC |
| **46** | ccattttaacggctagcatGCCAACTTTTCTTTTCTTTTTTGGGCCCATCCTGCAGGCTG |
| **47** | atgctagccgttaaaatggcttctaac |
| **48** | ATCCCGTCTAGAATCCGCgtag |
| **49** | TTCTAGACGGGATCCACCGGTCGCCACCATGgtgagcaagggcgaggag |
| **50** | gattatgatctagagtcgcggccgctcgagTTACTTGTACAGCTCGTC |

**Supplementary Figure 1**

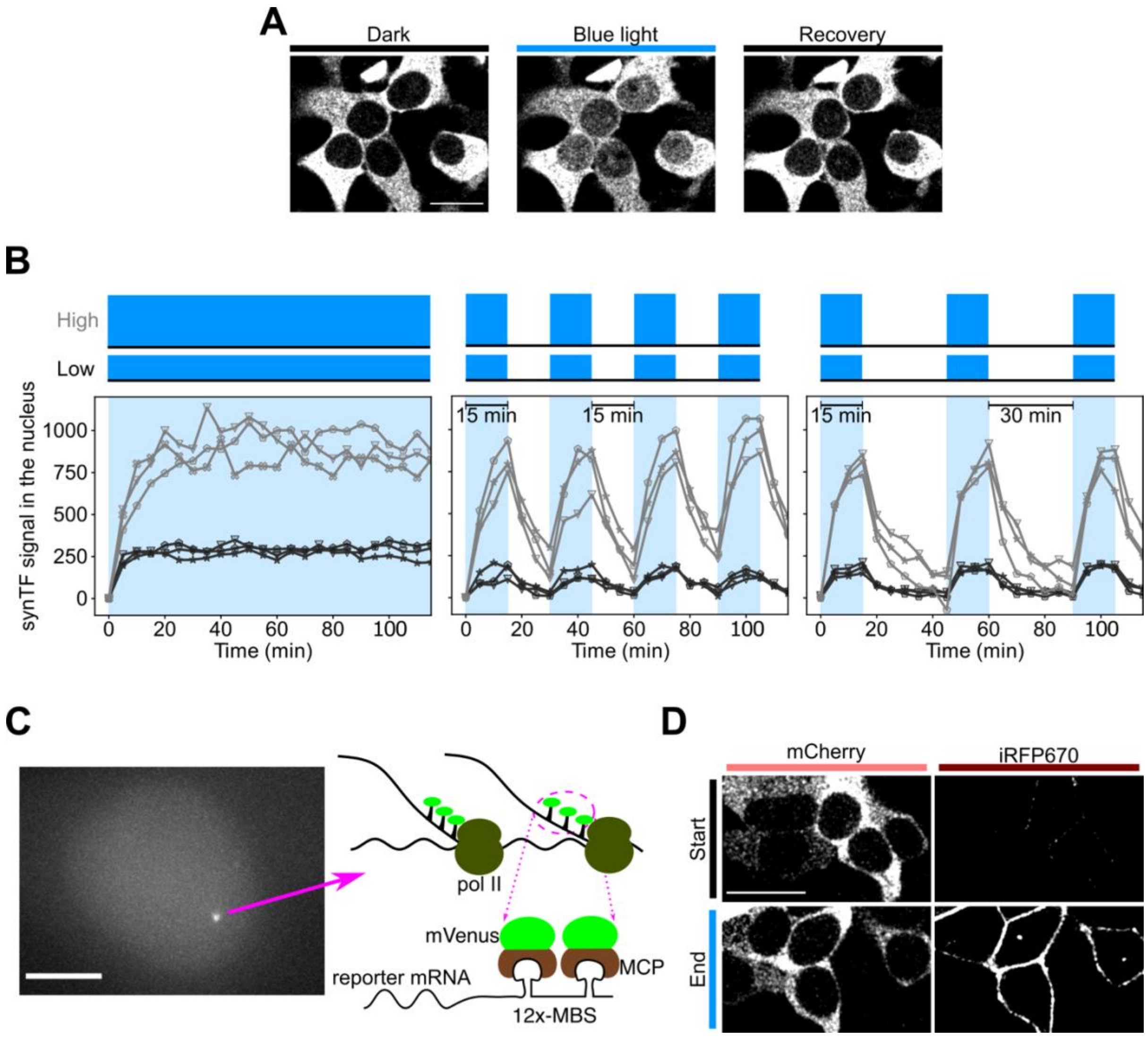


**Figure S1, related to Figure 1. Design of molecular components and experimental setup used to study how transcription factor dynamics are decoded by mammalian gene regulatory elements.**

(A) Representative fluorescence microscopy images showing accumulation of synTF in the nucleus of HEK293 cells upon blue light activation and recovery of cytoplasmic localization when the cells are kept in the dark. Illumination was performed shining 6.44 $Wm^{-2}$ blue light for 125 msec every 45 sec for 15 min. Scale bar, 20 μm.

(B) Generation of different TF dynamics with synTF. Graphs showing the nuclear TF signal over time in three distinct cells. Grey curves, high amplitude signal. Black curves, low amplitude signal. The low amplitude signal was achieved by illuminating the cells every 90 sec instead of every 45 sec.

(C) Setup for live cell imaging of nascent RNA. To monitor nascent RNAs in living cells, we deployed the MS2/MCP system (*55*, *56*). This comprises the bacteriophage MS2 capsid protein

(MCP), fluorescently labeled by means of a genetic fusion to an FP (mVenus in our case), and multiple repeats of sequence-specific RNA stem loops (twelve in our case), which are integrated in the reporter transcript at the 5’ or 3’ UTR (3’ UTR in our case). The stem loops are specifically bound by MCP, rendering the transcripts visible under the microscope at the site of transcription. Left, representative fluorescence microscopy image showing a fluorescent focus, which indicates the presence of several nascent RNAs. Scale bar, 5 μm. Right, schematic representation of the RNA visualization method used. MCP, MS2 coat protein. MBS, MCP binding site. MCP is expressed as a fusion to mVenus. Twelve repeats of the MCP binding sites (also called MS2 loops) are cloned in the 3’ UTR of the reporter.

(D) Representative fluorescence microscopy images of HEK293 cells showing synTF (mCherry) and reporter (iRFP670) levels before and after illumination with blue light. The reporter protein, iRFP670, is fused to the CAAX motif for plasma membrane localization. Scale bar, 20 μm.

**Supplementary Figure 2**

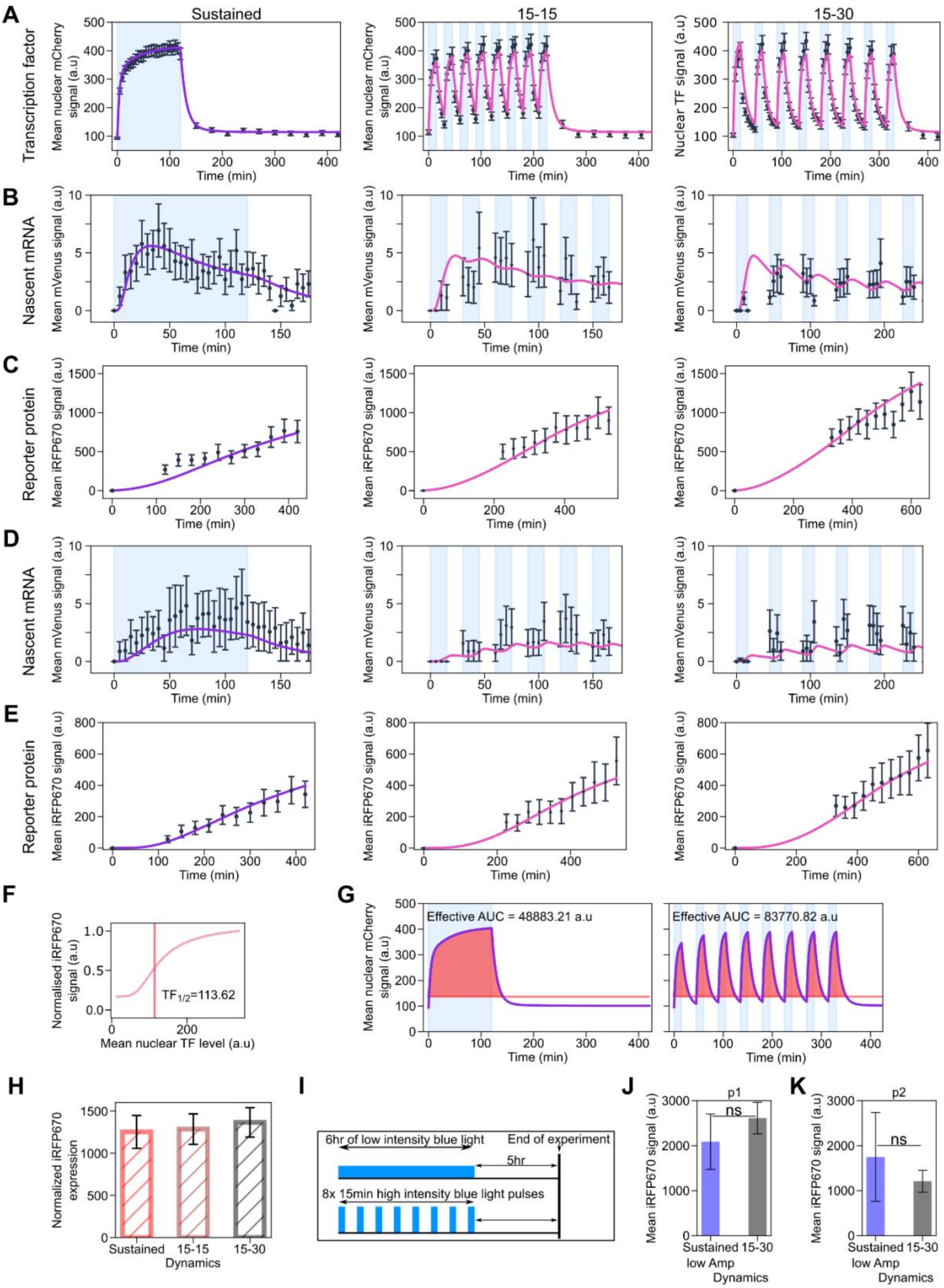


**Figure S2, related to Figure 2. Characterization of promoters p1, p2 and p3.**

(A) Quantification of synTF nuclear translocation over time for the indicated TF dynamics.
(B,D) Quantification of mean reporter nascent RNA over time for the indicated TF dynamics.
(C,E) Quantification of mean reporter protein levels over time for the indicated TF dynamics.
(A-C) The promoter used in these experiment was p2.
(D-E) The promoter used in these experiment was p3.
(F) Prediction of synTF concentration above which the reporter protein is expressed above the half maximal value.
(G) Calculation of the effective synTF cumulative levels using the threshold calculated in (F) for sustained (upper panel) and 15-30 pulsatile (lower panel) dynamics.
(H) Quantification of mean reporter protein levels at the end of the experiment for the indicated synTF dynamics normalized using the effective cumulative synTF levels calculated in (G).
(F-G) The promoter used was p1.
(I) Schematic showing the experimental setup in which amplitude was varied to achieve similar cumulative synTF levels at fixed experimental time.
(J,K) Quantification of mean reporter protein levels for the indicated TF dynamics for promoters p1 (J) and p2 (K). Light blue shadowing, blue light illumination phase. Together with the experimental data (black dots), fitted (violet line; for sustained dynamics) and simulated (pink line; for both pulsatile dynamics) values are shown. The mathematical model is shown in Figure 4 and the equations are described in the Supplememtary Text.
(A-E, H, J-K) Data represent mean ± s.e.m. of at least n=20 individual cells, imaged on at least n=3 biologically independent experiments. P-values were calculated with the Welch's t-test. ns, non signficant (P > 0.05).

**Supplementary Figure 3**

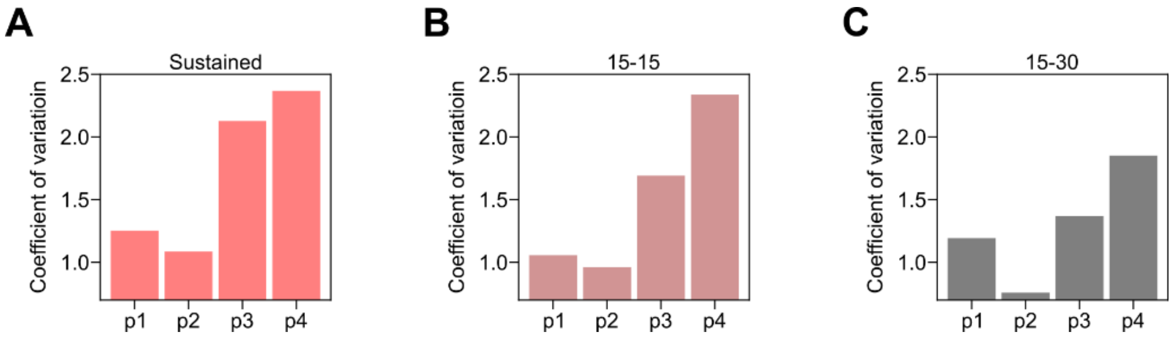


**Figure S3, related to Figure 3. Promoters p3 and p4 are noisy.**
(A-C) Quantification of the coefficient of variation for the mean reporter protein levels at the end of the experiments for the indicated promoters under sustained (A), 15-15 (B) and 15-30 (C) pulsatile dynamics.

**Supplementary Figure 4**

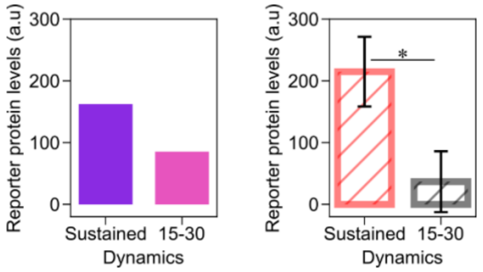


**Figure S4, related to Figure 5. Promoter p2 with two instead of four REs senses dynamics also at the protein level.**

Model predictions (left panel; fitted (violet bar) and simulated (pink bar) and experimental data (right panel) for the mean reporter protein levels at the end of the experiment for the indicated synTF dynamics in combination with promoter p8, which is a version of promoter p2 with two insteaf of four REs. Data represent mean ± s.e.m. of at least n=20 individual cells, imaged on at least n=3 biologically independent experiments. P-values were calculated with the Welch's t-test. *, P-value = 0.01545.

**Supplementary Figure 5**

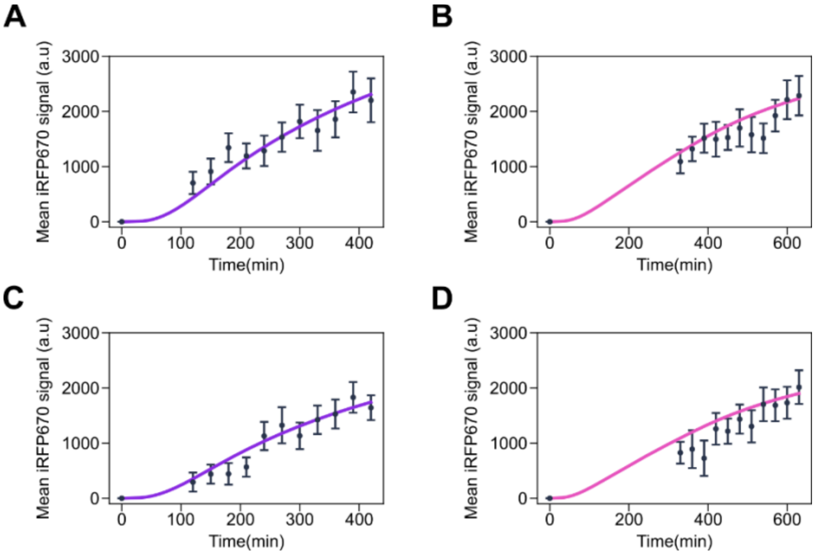


**Figure S5, related to Figure 5. Measurements of mean reporter protein levels over time for constructs 5UTR1 and 5UTR2.**

(A-D) Quantification of mean reporter protein levels over time for sustained (A,C) and 15-30 pulsatile (B,D) dynamics for constructs 5UTR1 (A,B) and 5UTR2 (C,D). Data represent mean ± s.e.m. of at least n=20 individual cells, imaged on at least n=3 biologically independent experiments. Lines represent simulations. Violet lines represent data fitting, while pink lines represent predictions.

**Supplementary Video 1**

**Video S1. Nascent RNA visualization for promoter p1 under sustained synTF dynamics.** HEK293 cells expressing NLS-MCP-mVenus for the visualization of the target RNA were transiently transfected with the plasmid encoding synTF and the reporter gene iRFP670 under promoter p1. synTF was subjected to sustained dynamics. Nascent RNA was visualized every 5 min for two hours.
